# Characterization of Lysine Methylation During Neuronal Differentiation of LUHMES cells

**DOI:** 10.64898/2025.12.31.696910

**Authors:** Jocelyne N. Hanquier, Malini Iyer, Taylor N. Evans, Devon L. McCourry, Christine A. Berryhill, Emma H. Doud, Whitney R. Smith-Kinnaman, Amber L. Mosley, Evan M. Cornett

## Abstract

Over one-third of human lysine methyltransferases (KMTs) and lysine demethylases (KDMs)-the enzymes responsible for adding or removing methylation on lysine residues within proteins-are linked to neurodevelopmental disorders (NDDs). Consequently, several studies have explored the roles of specific KMTs or KDMs in neuronal differentiation, and alterations in histone methylation patterns have been identified. It is now widely recognized that KMTs and KDMs also target non-histone proteins, yet knowledge of how non-histone lysine methylation changes during neuronal differentiation remains limited. Here, we address this gap using quantitative mass spectrometry-based proteomics to identify and measure changes in non-histone lysine methylation at three different stages in the Lund human mesencephalic (LUHMES) neuronal differentiation model. We identify 74 lysine methylation sites with significant differences in abundance across differentiation. Our analysis reveals lysine methylation on many non-histone proteins involved in neuronal differentiation and neurodevelopment, including signaling molecules, cytoskeletal proteins, RNA splicing factors, and transcription factors. Overall, this work broadens the understanding of non-histone lysine methylation in a neuronal differentiation model and offers a valuable resource of lysine methylation sites on proteins of biological and clinical significance for future research.

## Introduction

Lysine methylation (Kme) – a widespread, reversible post-translational modification (PTM) of histone and non-histone proteins – regulates diverse molecular functions and cellular processes, with effects that vary depending on methylation state and cellular context. The epsilon amine group of the lysine side chain can receive up to three methyl moieties, resulting in mono-(Kme1), di-(Kme2), or tri-methylation (Kme3). This reversible PTM is added by lysine methyltransferases (KMTs), removed by lysine demethylases (KDMs), and recognized by proteins containing methyllysine reader domains, triggering downstream molecular functions. Kme has been most extensively studied within the context of histone methylation, which has established roles in processes such as transcriptional activation and repression, heterochromatin formation, and genomic imprinting^1^. Nonetheless, more recent research efforts focusing on non-histone lysine methylation have resulted in the identification of over 10,000 Kme sites on close to 5,000 human proteins^2^. While the functional annotation of non-histone Kme sites remains limited, non-histone lysine methylation has been associated with molecular functions including protein-protein interactions, protein-DNA interactions, protein activity, protein stability, PTM crosstalk, and subcellular localization^3^.

KMTs and KDMs are critical for proper brain development, and dysregulation of these enzymes has critical developmental and cellular consequences^1^. Over a third of KMTs/KDMs have been associated with neurodevelopmental disorders (NDDs) through sequencing and functional studies^4–6^. Haploinsufficiency of a number of these NDD-associated enzymes results in aberrations in neuronal differentiation and development^7–9^, suggesting a role for Kme in these processes. As KMTs and KDMs are differentially expressed in models of embryonic brain development and human neuronal differentiation^10,11^, it stands to reason that the Kme events mediated by these enzymes may also change in abundance across differentiation. Lysine methylation of histone proteins has been demonstrated to regulate expression of genes critical for neuronal function and differentiation, including genes involved in synaptic formation, axogenesis, and pluripotency^12^. Less is understood about the functional role of non-histone Kme in neuronal differentiation due to limited reports; however, lysine methylation of tubulin and actin, two cytoskeletal proteins, has been demonstrated to have critical roles in neuronal cell migration and polarization through regulation of microtubule dynamics^13,14^. While site-specific changes in histone methylation have been reported across neuronal differentiation^12^, to our knowledge, high-throughput profiling of global Kme across neuronal differentiation has not been conducted, precluding our understanding of the molecular mechanisms by which non-histone Kme regulates neuronal differentiation.

In this study, we sought to quantitatively profile global lysine methylation across differentiation using the Lund human mesencephalic (LUHMES) cells, which have previously been used to model neurodegenerative and neurodevelopmental disorders^15,16^. We employed Tandem Mass Tag (TMT) mass spectrometry (MS) to simultaneously profile global protein abundance and lysine methylation sites across three stages of LUHMES differentiation: the neural progenitor stage (differentiation day 0), an early differentiation stage (differentiation day 2), and the post-mitotic, dopaminergic-like neuron stage (differentiation day 6). Our study identified and quantified hundreds of Kme sites on diverse, non-histone proteins, revealing differentiation-stage-specific lysine methylome signatures. Overall, our results expand the scope of non-histone Kme events reported across neuronal differentiation.

## Materials and Methods

### LUHMES maintenance and differentiation

The LUHMES cell line used in this study was purchased from ATCC (CRL-2927). LUHMES cells were cultured and differentiated as previously described^17^. Culture dishes were pre-coated with 15 µg/mL poly-L-ornithine (MilliporeSigma #P4957) followed by 10 µg/mL laminin (MilliporeSigma #L2020) to promote cell adherence. LUHMES cells were cultured at 37°C with 5% CO_2_. Proliferating cells were maintained using complete medium containing: DMEM/F-12 (Corning #10-092-CV), 1% N2 Supplement (Fisher Scientific #17-502-001), and 40 ng/mL bFGF (MilliporeSigma #GF003AF). One day prior to differentiation, 3 million cells were freshly seeded into a new T75 flask in complete medium. The following day (differentiation day 0), complete medium was removed and cells were washed twice with PBS prior to adding differentiation medium: DMEM/F-12(Corning #10-092-CV), 1% N2 Supplement (Fisher Scientific #17-502-001, 1 mM cAMP (MedChemExpress #HY-B1511), 1 µg/mL doxycycline (MilliporeSigma #D5207), 2 ng/mL GDNF (MilliporeSigma #G1777), and 2 mM glutamax (ThermoFisher Scientific #35050061). On day 2 of differentiation, cells were removed from the flask using trypsin and 8 million cells were seeded into new T75 flasks in differentiation medium. On differentiation day 4, the differentiation medium was replenished. Cells were collected for mass spectrometry analysis on differentiation days 0, 2, and 6 (n=4 biological replicates per time point).

### Immunocytochemistry

Immunocytochemistry was performed as previously described^18^. Briefly, cells differentiated on 25 mm-diameter glass coverslips (Neuvitro #GG-25-1.5-PRE) were fixed with 4% paraformaldehyde for 20 minutes, then washed 3 times with PBS. Cells were permeabilized with 2% bovine serum albumin (BSA) and 0.3% Triton X-100 in PBS for 30 minutes, followed by 2 washes with PBS containing 2% BSA. Cells were then incubated with primary antibodies, Beta 3-tubulin (Cell Signaling Technology #4466S; 1:500 dilution) and MAP2 (Cell Signaling Technology #4542S; 1:1,000 dilution), in PBS containing 2% BSA overnight at 4°C. Following primary antibody incubation, cells were washed for 3 times for 5 minutes with PBS and incubated with fluorescent secondary antibodies, either goat anti-rabbit 647 (Invitrogen #A32733) or goat anti-mouse 488 (Invitrogen #A32723), in PBS containing 2% BSA for 1 hour at room temperature. Following secondary antibody incubation, cells were washed 3 times with PBS for 5 minutes each. Coverslips were then mounted to glass slides using ProLong Gold Antifade with DAPI (ThermoFisher #P36930) and cured for 24 hours at room temperature in the dark. Slides were imaged using a Zeiss AxioObserverZ1 modified by 3i (www.intelligent-imaging.com) for confocal microscopy, and images were analyzed using ImageJ/Fiji Bio-Formats plugins^19^. Within any set of comparable images, the image capture and scaling conditions were identical. The cyan/blue channel included a 405-emission filter (DAPI), the green channel included a 488**-**emission filter (Beta 3-tubulin), and the red channel included a 560-emission filter (MAP2). Z-stack imaging was performed, and a 5-layer stack was generated using Fiji to enhance visualization. The Z-projection of the stacked images was set to “max intensity,” and brightness and contrast were adjusted for each channel using designated maximum and minimum intensity levels to optimize visualization.

### Western blotting

Cells were grown to 80% confluence and resuspended in lysis buffer: 1X RIPA buffer (Cell Signaling Technology #9806) supplemented with protease inhibitors (Pierce #A32953) and endonuclease (Pierce # 88700). Lysates were run on 8% SDS-PAGE gels and probed with anti-mono methyl lysine (Kme1) (Cell Signaling Technology #14679S; 1:1,000 dilution), anti-di-methyl lysine (Kme2) (Cell Signaling Technology #14117S; 1:1,000 dilution), or anti-tri-methyl lysine (Kme3) (PTM Bio #PTM-601; 1:1,000 dilution). Ponceau red staining of the membrane was used to visualize total protein prior to antibody probing.

### Quantitative Proteomics Sample Preparation and Labeling

Cells were collected for mass spectrometry analysis on differentiation days 0, 2, and 6 (n=4 biological replicates per time point). Cell pellets were resuspended in 400 µL 8 M Urea, 100 mM Tris pH 8.5 and transferred to Diagenode Bioruptor tubes (#C30010010). Cells were lysed via Diagenode Bioruptor, 30s on/30s off, for 30 cycles. Samples were then clarified, and protein concentration determined by Bradford assay(Biorad #500201). 25 µg of each sample was treated with 5 mM Tris(2-carboxyethyl)phosphine hydrochloride (TCEP, Sigma-Aldrich #C4706) to reduce disulfide bonds and the resulting free cysteine thiols were alkylated with 10 mM chloroacetamide (CAA, Sigma Aldrich #C0267). Samples were diluted with 50 mM Tris.HCl pH 8.5 (Sigma-Aldrich #10812846001) to a final urea concentration of 2 M for overnight Trypsin/Lys-C digestion at 35 °C (1:50 protease: substrate ratio, Mass Spectrometry grade, Promega Corporation #V5072). In addition to the 12 samples that were digested with Trypsin/LysC, 12 samples were reduced and alkylated as stated above, then diluted to <0.5 M Urea with 100 mM Tris-HCl and 10 mM CaCl_2_, pH 7.8, and digested with chymotrypsin overnight at 35 °C (1:25 protease: substrate, Roche #11418467001).

Digestions were quenched with trifluoroacetic acid (TFA, 0.5% v/v) and desalted on Waters Sep-Pak® Vac cartridges (Waters™ #WAT054955) with a wash of 1 mL 0.1% TFA followed by elution in 0.6 mL of 70% acetonitrile 0.1% formic acid (FA). Peptides were dried by speed vacuum and resuspended in 50 mM triethylammonium bicarbonate. Samples were divided into two 12-plex samples. Each sample was then labeled for overnight at room temperature, with 0.5 mg of Tandem Mass Tag Pro (TMTpro) reagent (16-plex kit, manufacturer’s instructions, Thermo Fisher Scientific, TMTpro™ Isobaric Label Reagent Set; # 44520, lot #ZA382395). Prior to quenching, labeling efficiency of over 95% was confirmed.

Reactions were quenched with 0.3 % hydroxylamine (v/v) at room temperature for 15 minutes. Labeled peptides were then mixed and dried by speed vacuum. The Chymotrypsin 12-plex was resuspended in 0.5% TFA and fractionated on a Waters Sep-Pak® Vac cartridge (Waters™ #WAT054955) with a 1 mL wash of water, 1 mL wash of 5% acetonitrile, 0.1% triethylamine (TEA) followed by elution in 12.5%, 15%, 17.5%, 20%, 22.5%, 25%, 30%, and 70% acetonitrile, all with 0.1% TEA). The Trypsin/Lys-C 12-plex was resuspended in 10 mM ammonium formate, pH 10. Half of each mix was fractionated using an offline Thermo UltiMate 3000 HPLC with a Waters Xbridge C18 column (3.5 µm x 4.6mm x 250 mm, Cat No: 186003943; Buffer A: 10 mM formate pH 10, Buffer B: 10 mM formate pH 10, 95% acetonitrile, gradient 1 mL/min 0-15%B over 5 min, 15-20% B over 5 min, 20-35%B over 75 min, 35-50% B over 5 min, 50-60% B over 10 min and a 6 minute hold at 60% B). Fractions were collected continuously every 60 seconds into 9-well plates. Initial and late fractions with minimal material were combined and lyophilized. The remaining fractions were concatenated into 24 fractions, dried down, and resuspended in 50 µL 0.1% FA prior to online LC-MS^20,21^.

Mass spectrometry was performed utilizing an EASY-nLC 1200 HPLC system (SCR: 014993, Thermo Fisher Scientific) coupled to Orbitrap Fusion™ Eclipse™ mass spectrometer (Thermo Fisher Scientific) with FAIMSpro interface (Thermo Fisher Scientific). Each multiplex was run on an Aurora column with 180 minute gradient. For each fraction, 25% of the sample was loaded onto an Aurora column, and run at 350 nl/min. (Thermo Fisher ES902). The gradient (Mobile phases A: 0.1% formic acid (FA), water; B: 0.1% FA, 80% Acetonitrile (Thermo Fisher Scientific Cat No: LS122500)), was increased from 8-38%B over 160 minutes; 95% B over 12 mins; and dropping from 95%-5% B over the final 8 min. The mass spectrometer was operated in positive ion mode, default charge state of 2, advanced peak determination on, and Easy IC™ on. Three FAIMS compensation voltages (CVs) were utilized (-45 CV; -55 CV; -65CV) each with a cycle time of 1 s and with identical MS and MS2 parameters. Precursor scans (m/z 400-1600) were done with an orbitrap resolution of 120000, RF lens% 30, dynamic maximum inject time, standard AGC target, minimum MS2 intensity threshold of 2.5 e4, MIPS mode to peptide, including charges of 2 to 7 for fragmentation with 60 sec dynamic exclusion. MS2 scans were performed with a quadrupole isolation window of 0.7 m/z, 34% HCD CE, 15000 resolution, standard AGC target, automatic maximum IT, and fixed first mass of 100 m/z.

### Data analysis

The resulting RAW files were analyzed in Proteome Discover™ 2.5 (Thermo Fisher Scientific) using a *Homo sapiens* UniProt-reviewed proteome FASTA (downloaded 051322, 20282 sequences) and common laboratory contaminants (73 sequences). SEQUEST HT searches were conducted with full trypsin digest, a maximum of 5 missed cleavages, precursor mass tolerance of 10 ppm, and a fragment mass tolerance of 0.02 Da. Static modifications used for the search were: 1) carbamidomethylation on cysteine (C) residues; 2) TMTpro label on N-termini of peptides. Dynamic modifications used for the search were 1) TMTpro label on lysine (K) residues, 2) methylation on lysine (K) residues, 3) dimethylation on lysine (K) residues, 4) trimethylation on lysine (K) residues, 5) oxidation of methionine, 6) methionine loss, or 7) acetylation with methionine loss on protein N-termini. A maximum of 5 PTMs was allowed per peptide. The same parameters were used for the chymotrypsin digest, except for the use of chymotrypsin as the protease. Percolator False Discovery Rate was set to a strict setting of 0.01 and a relaxed setting of 0.05. The IMP-ptm-RS node was used for all modification-site localization scores. Values from both unique and razor peptides were used for quantification. In the consensus workflows, peptides were normalized to the total peptide amount without scaling. Quantification methods utilized TMTpro isotopic impurity levels available from Thermo Fisher Scientific. Reporter ion quantification used a minimum S/N threshold of 6 and a co-isolation threshold of 30%. Resulting grouped abundance values for each sample type, abundance ratio (AR) values, and respective p-values (t-test) from Proteome Discover were exported for further analysis.

All downstream data analysis was performed using R version 4.4.0. Proteins were filtered for a high protein FDR confidence with two or more unique peptides, and a pseudo count of 0. 1 was added to the normalized abundances to circumvent data analysis run errors. Lysine methylated peptides were filtered for a Site Localization Score (ptmRS module) greater than or equal to 0.9 and unambiguous. To account for changes in protein abundance across differentiation, the Kme peptide-normalized abundance values were normalized to the protein-normalized abundance values for each biological replicate. Differentially abundant proteins and protein-normalized Kme peptides were determined using an ANOVA (padj < 0.05) followed by Tukey pair-wise comparisons (padj<0.05). Data from PhosphoSitePlus®^22^ was accessed in January 2025, and data from InterPro^23^, Uniprot^24^, ClinVar^25^, and Simons Foundation Autism Research Initiative (SFARI) database^6^ was accessed in March 2025. The following R packages were used for the indicated analyses: *GGally* (Pearson correlation analysis), *ggplot2* (visualization of principal component analyses, motif analyses, and boxplots), *ggpubr* (adding statistics to boxplots), *ggvenn* (venn diagram visualization), *pheatmap* (heatmap visualization), *EnhancedVolcano* (volcano plot visualization), *UpSetR* (upset plot visualization). Histogram visualization was performed using the “hist” function from the base R package *graphics*. Protein and Kme peptide co-expression modules were identified using the *WGCNA* R package for weighted gene co-expression network analysis using a signed network approach^26^. A soft-threshold power of 30 (protein WGCNA) or 18 (Kme peptides) was used. Modules with a minimum size of 30 (protein) or 10 (Kme peptides) were identified using hierarchical clustering.

## Results

### Quantitative profiling of the proteome and lysine methylome across LUHMES differentiation

To characterize stage-specific changes in the proteome and lysine methylome of LUHMES cells across differentiation, quantitative mass spectrometry analysis was conducted on cells spanning three distinct stages of LUHMES differentiation. LUHMES human neural progenitor cells were differentiated into post-mitotic, dopaminergic-like neurons using an established protocol^17^. Immortalized via doxycycline-controlled v-myc overexpression, LUHMES were differentiated within six days following exposure to doxycycline, glial-derived neurotrophic factor (GDNF), and dibutyryl (db)-cAMP (**Figure 1A**). Differentiated LUHMES cells displayed morphological characteristics of neurons, including neurite projections (**Figure 1A**), and expressed neuronal maturation markers, including β3-tubulin and MAP2 (**Figure S1A**). Changes in lysine methylation of non-histone proteins across LUHMES differentiation were detected *in vitro* by immunoblotting using pan-methyl lysine antibodies against all three methyl states (mono-, di-, and tri-methylation) (**Figure S1B**). The most pronounced changes in lysine methylation were detected using a pan-tri-methyl antibody. Most methylated proteins detected decreased in abundance across differentiation; however, this could be due to the sequence selectivity of pan-methyl lysine antibodies and the detection of methylation on the most highly abundant proteins^27^.

**Figure 1:**
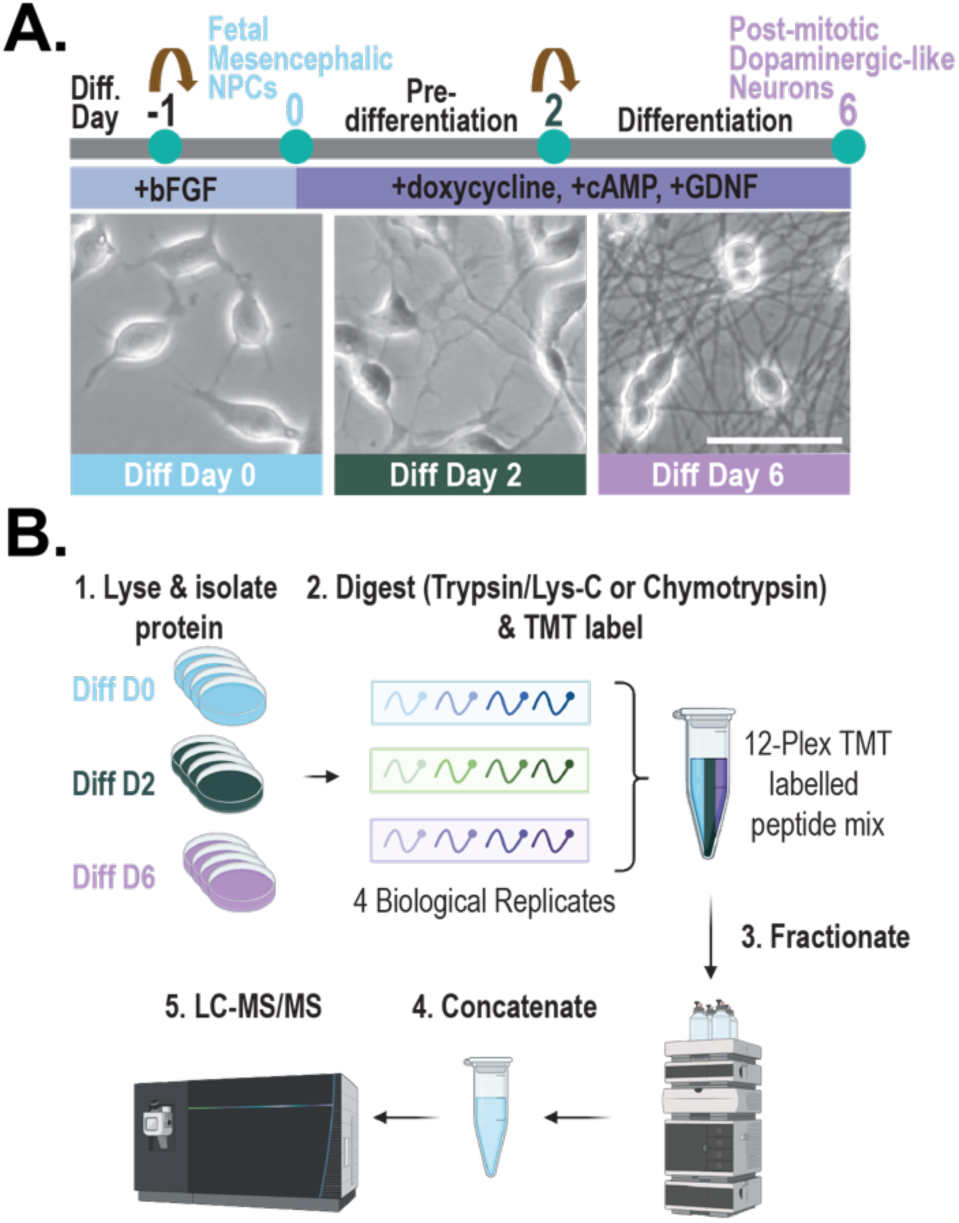
Experimental approach for quantitative profiling of the LUHMES proteome and lysine methylome. **(A)** Schematic diagram of Lund human mesencephalic (LUHMES) neural progenitor cell differentiation into post-mitotic, dopaminergic-like neurons following 6-day exposure to doxycycline, glial-derived neurotrophic factor (GDNF), and dibutyryl (db)-cyclic adenosine monophosphate (cAMP). Brightfield microscopy images of LUHMES cells. Scale bar represents 50 µm. **(B)** Quantitative proteomics approach for profiling the LUHMES proteome and lysine methylome. LUHMES cells (differentiation days 0, 2, or 6; n=4) were collected, lysed, and trypsin/Lys-C or chymotrypsin digested into peptides. Each biological replicate received an isobaric tandem mass tag (TMT) chemical label, resulting in two, 12-plex TMT labelled peptide mixtures (one for tryptic/Lys-C and one for chymotryptic peptides). Each of the 12-plex TMT labelled peptide mixtures was subjected to high pH fractionation followed by tandem mass spectrometry (LC-MS/MS) analysis, resulting in two MS experiments. Proteome Discoverer (2.5) was used for database searching.

To quantitatively profile changes in protein and lysine methylated peptide abundance across LUHMES differentiation, we employed the experimental approach as follows: LUHMES cell lysates (n=4) were collected on differentiation days 0 (neural progenitor stage), 2 (early stage of differentiation), and 6 (post-mitotic, dopaminergic-like neuron stage) and digested into peptides using either trypsin/lys-C or chymotrypsin. Peptides from each biological replicate were labeled with a unique isobaric tandem mass tag (TMT), resulting in two, 12-plex TMT-labeled peptide mixtures (one for tryptic/lys-C and one for chymotryptic peptides). Each of the 12-plex TMT labelled peptide mixtures was subjected to off-line fractionation followed by tandem mass spectrometry (LC-MS/MS) analysis, resulting in two MS experiments (**Figure 1B**). In the MS experiment with trypsin/lys-C digestion, 146,016 peptide spectral matches (PSMs), 56,492 peptides, 5,516 detected proteins, and 5,428 quantified proteins were identified. In the MS experiment with chymotrypsin digestion, 109,777 PSMs, 25,779 peptides, 2,800 detected proteins, and 2,755 quantified proteins were identified. In total, 5,655 proteins were detected, and 5,568 proteins were quantified. As expected, more proteins were detected following trypsin/lys-C digestion compared to chymotrypsin digestion. This is likely due to the properties of tryptic peptides, including basic residues at the N- and C-termini, that render them favorable for protonation and ionization, triggering detection by MS^28^. Nonetheless, approximately 95% (n= 2,615) of all proteins quantified in the chymotrypsin MS experiment were also quantified in the trypsin/lys-C experiment (**Figure S2**).

Pearson linear correlation analysis of protein abundance reveals high reproducibility within and between the two MS experiments. The protein abundance of the biological replicates demonstrated a high degree of reproducibility within the trypsin/lys-C (n = 5,428; R ≥ 0.993; p < 0.05) and chymotrypsin (n = 2,755; R ≥ 0.957; p < 0.05) experiments (**Figure S3A**). Principal component analysis (PCA) of the protein abundances revealed distinct clusters of biological replicates for each stage of differentiation across both experiments (**Figure S3B**). Furthermore, Pearson correlation analysis of the average protein abundances on differentiation days 2 and 6 relative to day 0 revealed a high degree of reproducibility between the two experiments (n = 2,615; R ≥ 0.876; p < 0.05) (**Figure S4**). Additionally, protein abundance changes quantified in this study correlate with protein abundance changes from a study conducted by Tüshaus et al.^10^, in which undifferentiated and day 6 differentiated LUHMES cells were subjected to MS analysis and label-free quantification. Approximately 85% (4,551 of 5,344) of the proteins quantified in the Tüshaus study were also quantified between the two MS experiments in this study (**Figure S5A**), and Pearson correlation analysis of the averaged protein abundances on differentiation day 6 relative to day 0 revealed a high degree of reproducibility between the shared proteins quantified in the Tüshaus study and those quantified in both our trypsin/lys-C and chymotrypsin experiments (n=2,551; R>0.757; p<0.05) (**Figures S5B-C**).

### Stage-specific changes in protein abundance of differentiation markers across LUHMES differentiation

In this study, hundreds (chymotrypsin MS) to thousands (trypsin/lys-C MS) of proteins were determined to be differentially abundant across distinct stages of LUHMES differentiation (ANOVA padj<0.05 followed by Tukey pairwise padj<0.05) (**Figure S6**). To identify clusters of proteins with similar expression patterns across differentiation, weighted gene correlation network analysis (WGCNA)^26^ was performed. WGCNA analysis revealed 7 protein co-expression modules (plus a ‘gray’ module comprised of proteins not assigned to a co-expression module) with distinct expression patterns across differentiation (**Figures 2A and S7A-B**). Gene ontology (GO) enrichment analysis of the WGCNA modules revealed significant enrichment for different biological processes across protein modules (**Figures 2B and S7C**).

**Figure 2:**
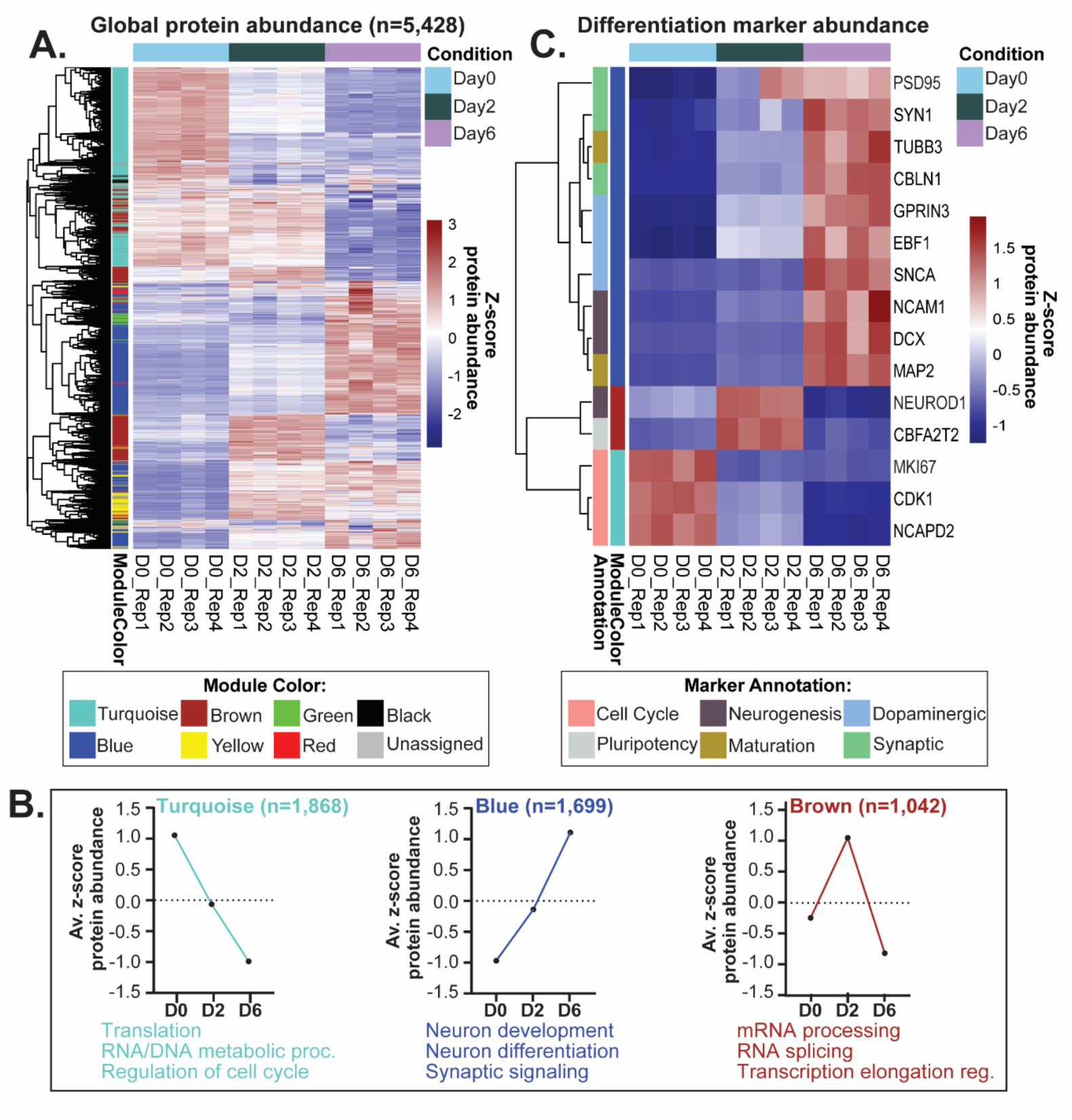
Stage-specific changes in protein abundance across LUHMES differentiation. **(A)** Heatmap of the proteins (n=5,428) quantified across LUHMES differentiation in the trypsin-/Lys-C experiment. Colors represent the z-score of protein abundance. Rows (proteins) and columns (differentiation sample) are clustered by Euclidian distance. The co-expression color module associated with a given protein, through weighted gene correlation network analysis (WGCNA), is represented to the left of the rows. **(B)** Expression profiles (z-score protein abundance) and enriched gene ontology (GO) biological processes (p<0.05) corresponding to the indicated WGCNA modules. **(C)** Heatmap of protein abundance of a subset of differentiation markers corresponding to the indicated WGCNA modules. Rows (proteins) and columns (differentiation sample) are clustered by Euclidian distance. Functional annotation of protein represented to the left of the rows. Names of genes that encode the proteins are displayed to the right of the rows.

As expected, protein modules are comprised of established differentiation markers, as well as module-specific proteins of interest (**Figure 2C**), and expression changes are congruent with those reported across differentiation of human midbrain dopaminergic neurons^11^. For example, proteins that regulate cell cycle progression and pluripotency were more highly expressed in early stages of LUHMES differentiation (turquoise and brown modules, respectively), and proteins that are associated with neuronal maturation, dopaminergic activity, and synaptic activity were more highly abundant at the neuronal stage of differentiation (blue module).

The turquoise module (n=1,868 proteins) comprises proteins that decreased in abundance during differentiation. Within this module, there is an enrichment for proteins involved in translation, RNA/DNA metabolic processes, and regulation of the cell cycle. Proteins associated with the turquoise module include established cell cycle markers, such as Cyclin-dependent kinase 1 (*CDK1*) and Proliferation marker protein Ki-67 (*MKI67*), as well as proteins involved in chromosome condensation, such as Condensin complex subunit 1 (*NCAPD2*), which functions as a regulatory subunit of the condensin complex required for mitotic chromosome condensation, a critical process regulating mammalian cerebral cortex size^29^.

The brown module (n=1,042 proteins) comprises proteins that increased in abundance from differentiation day 0 to day 2, then decreased from day 2 to day 6. Within this module, there is an enrichment for proteins involved in mRNA processing, RNA splicing, and transcription elongation regulation. Proteins associated with the brown module include established neurogenesis markers, such as the pro-neural transcription factor Neurogenic differentiation factor 1 (*NEUROD1*). Single-cell RNA-sequencing data previously found that *NEUROD1* expression is enriched in human neural progenitors and neuroblasts across ventral midbrain development^11^. Another protein associated with the brown module is Protein CBFA2T2 (*CBFA2T2*), a transcriptional co-repressor that acts synergistically with other transcription factors, including the germline-specific transcription factor PRDM14, to regulate pluripotency and germline specification in mice^30^.

The blue module (n=1,699 proteins) comprises proteins that increase in abundance across differentiation. Within this module, there is an enrichment for proteins involved in neuron development, neuron differentiation, and synaptic signaling. Proteins associated with the blue module include established markers: neurogenesis markers [Neuronal migration protein doublecortin (*DCX*) and Neural cell adhesion molecule 1 (*NCAM1*)], maturation markers [Tubulin beta-3 chain (*TUBB3*) and Microtubule-associated protein 2 (*MAP2*)], dopaminergic markers [Alpha-synuclein (*SNCA*), G protein-regulated inducer of neurite outgrowth 3 (*GPRIN3*), and Transcription factor COE1 (*EBF1*)], and synaptic markers [Synapsin-1 (*SYN1*), Postsynaptic density protein 95 (*PSD95*), and Cerebellin-1 (*CBLN1*)]. As a synaptic organizer, Cerebellin-1 acts on pre- and post-synaptic components, is critical for synapse formation and maintenance^31^, and acts as a cue for axon growth and guidance throughout the central nervous system^32^. Taken together, the state-specific protein co-expression modules highlight biological processes that are critical for the differentiation of neural progenitor cells into dopaminergic-like neurons.

### Changes in protein abundance of lysine methyltransferases and demethylases across differentiation

In addition to markers of differentiation, a subset (38 out of ∼90) of the enzymes that mediate the addition and removal of lysine methylation – lysine methyltransferases (KMTs) and lysine demethylases (KDMs), respectively – were quantified in this study. In our trypsin/lys-C experiment, 38 KMTs/KDMs were quantified, 7 of which were also quantified in the chymotrypsin experiment. Additionally, all 21 KMTs/KDMs quantified in the Tüshaus et al. study^10^ were quantified in this study, and Pearson correlation analysis indicates correlation of KMT/KDM abundance between the two studies (n=21; Pearson correlation R=0.786, p<0.05) (**Figure S8**). Of the 38 KMTs/KDMs quantified in the trypsin/lys-C experiment, 35 are differentially abundant across differentiation (ANOVA padj<0.05 followed by Tukey pairwise padj<0.05), and unsupervised hierarchical clustering of KMT/KDM abundance reveals distinct clusters of expression across differentiation (**Figures 3A-C**).

**Figure 3:**
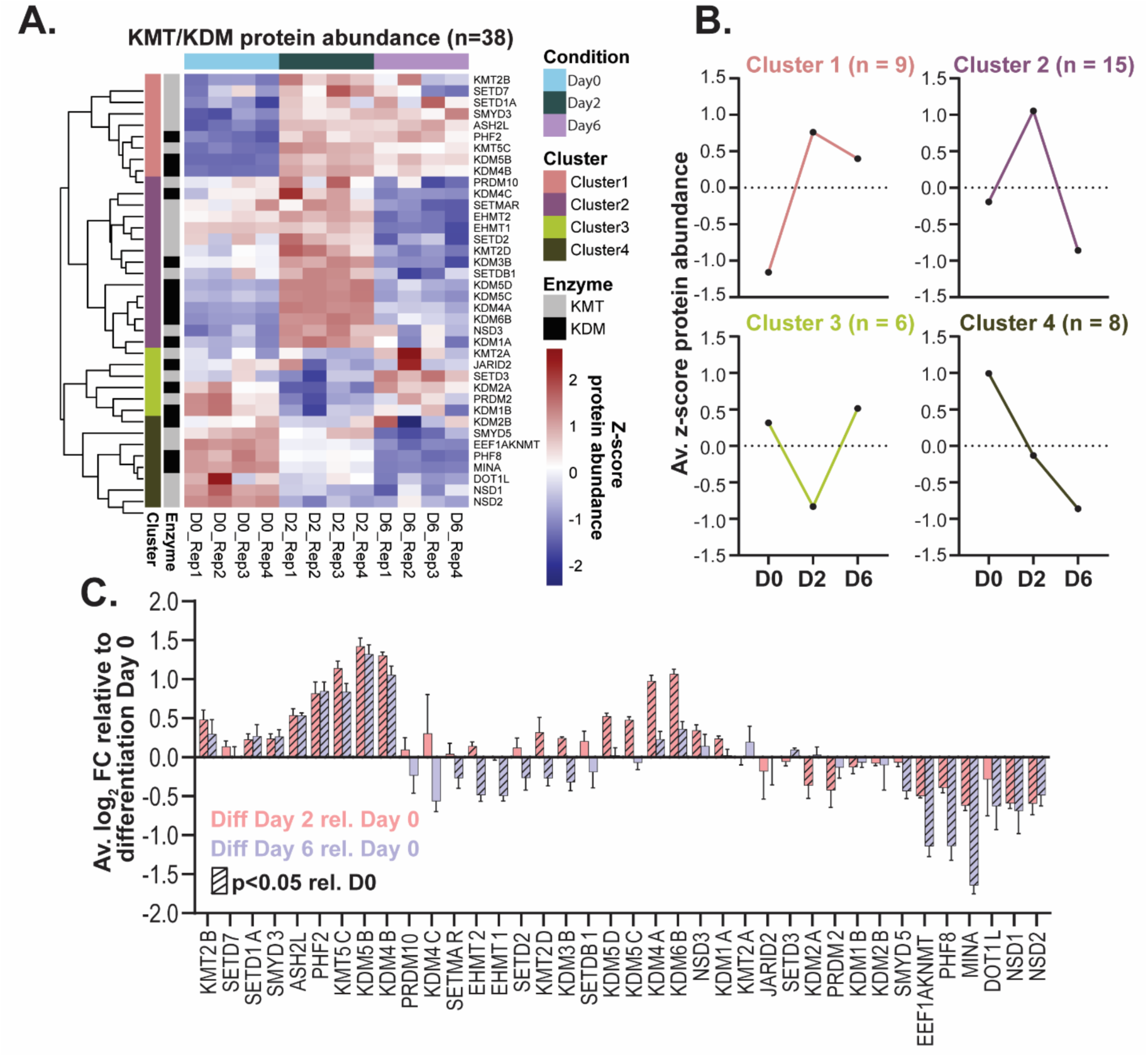
Changes in protein abundance of lysine methyltransferases (KMTs) and demethylases (KDMs) across LUHMES differentiation. **(A)** Heatmap of KMTs/KDMs (n=38) quantified across LUHMES differentiation in the trypsin-/Lys-C experiment. Colors represent the z-score of protein abundance. Rows (proteins) and columns (differentiation sample) are clustered by Euclidian distance. Enzyme classification (as a KMT or KDM) and enzyme expression cluster, revealed by hierarchical clustering, are represented to the left of the rows. Names of genes that encode the proteins are displayed to the right of the rows. **(B)** Expression profiles (z-score protein abundance) corresponding to clusters of KMT/KDM abundance are displayed. **(C)** Average log_2_ fold change of KMT/KDM abundance on differentiation days 2 (pink) and 6 (purple) relative to day 0. Diagonal stripes in bars indicate differential abundance (ANOVA padj<0.05 followed by Tukey pairwise padj<0.05) relative to day 0.

Several of the lysine methylation mediators quantified in this study, including the KMTs SMYD5 and EHMT2 (also known as G9a), play critical roles in regulating differentiation and have both histone and non-histone substrates. The decreased expression of SMYD5 and EHMT2 at the neuronal stage of LUHMES differentiation, relative to the neural progenitor stage (**Figure 3C**), is consistent with their reported roles in silencing of neuronal genes. SMYD5 knockdown in embryonic stem cells (ESCs) results in decreased ESC self-renewal and perturbed differentiation^33^. The mechanism associated with these phenotypes is loss of SMYD5-dependent silencing of heterochromatin through H4K20 tri-methylation, resulting in de-repression of differentiation genes. Additionally, there are reported functional roles for SMYD5 methylation of non-histone proteins; for instance, SMYD5-mediated tri-methylation of ribosomal protein RPL40 at K22 is associated with regulation of translation elongation^34^. Similarly, EHMT2 appears to be required for the maintenance of Polycomb repressive complex 2 (PRC2)-mediated gene silencing of a subset of developmental and neuronal genes in mouse ESCs. The proposed mechanism involves stimulation of PRC2 recruitment to common genomic loci via EHMT2-mediated mono-methylation of H3K27^35^. Additionally, EHMT2-mediated methylation of a variety of non-histone proteins, including Metastasis-associated protein MTA1, is associated with transcriptional repression^36^. The methylation status of MTA1 serves as a molecular switch for the formation of two protein complexes with opposite impacts on transcription regulation-the co-repressor nucleosome remodeling and histone deacetylase complex (NuRD) or the co-activator nucleosome-remodeling factor (NuRF) complex^37^. EHMT2-mediated mono-methylation of MTA1 at K532 is required for the formation and co-repressor activity of the NuRD complex^37^, which has been reported to regulate expression of genes critical for neuronal differentiation and brain development^38,39^. The characterization of KMT/KDM expression patterns is a critical initial step in determining the functions and lysine methylation substrates of these enzymes across different model systems.

### Detection of unique non-histone lysine methylation sites by MS following trypsin/lys-C and chymotrypsin digestion

We previously demonstrated that detection and quantification of hundreds of lysine methylation (Kme) sites across cell lines is possible using a similar TMT-based proteomics workflow, without immunoaffinity enrichment for lysine methylation, using trypsin/lys-C digestion^40^. In this study, we sought to determine the impact of different digestion strategies – trypsin/lys-C or chymotrypsin – on the detection of Kme sites by MS. Comprehensive *in vitro* biochemical characterization of KMT substrate selectivity using lysine-oriented peptide libraries, revealed that several KMTs prefer K- and R- rich sequences. While trypsin/lys-C digestion is the MS “gold standard”^41^, K- and R-rich peptides are challenging to detect after trypsin/lys-C digestion due to the peptide fragments being too small or the properties of the resulting peptides being incompatible for triggering MS detection^42^. Taken together, these observations led us to hypothesize that using alternative digestion strategies may lead to the detection of novel Kme sites.

In total, 303 methylated peptides corresponding to 273 unique Kme sites on 222 unique proteins were detected in this study (**Figures 4A-B**). Comparison of all Kme sites detected in this study to those reported in the PhosphoSitePlus® repository for PTMs (**Figure 4A**) and those detected in the Tüshaus study (based on our re-searching of the MS data for lysine methylation) (**Figure S9**) reveals that ∼90% of the detected Kme sites in this study are novel. Additionally, ∼50% of the unique proteins methylated in our study are not reported to be methylated in PhosphoSitePlus® (**Figure 4A**). Of the Kme sites detected in this study, ∼68% (n=185) are unique to the trypsin/lys-C experiment, ∼30% (n=82) are unique to the chymotrypsin experiment, and ∼2% (n=6) were detected in both experiments (**Figure 4B**). In addition to the minimal overlap in Kme sites detected, only ∼ 3% (n=6) of all unique methylated proteins were detected with both digestion strategies. We posited that digestion using chymotrypsin would result in the detection of more K- and R-rich sequences by MS compared to trypsin/lys-C. To test this hypothesis, analysis of the K and R density of Kme sites detected by either digestion strategy was performed. Comparison of 7-mer sequences surrounding methylated lysine residues reveals that, regardless of digestion strategy, ∼60% of all lysine-centered 7-mer sequences contain no additional K/R residue and ∼35% have one additional K/R residue, regardless of digestion strategy (**Figure S10A**). Comparison of the actual methylated peptide sequences detected by MS shows a slight shift in the K- and R-density of peptides detected between digestion strategies. Approximately 18% of peptide sequences detected following chymotrypsin digestion, compared to ∼8% of sequences detected following trypsin/lys-C digestion, contain a total of 4 K/R residues (**Figure S10B**). To evaluate the sequence context surrounding detected lysine methylation sites, motif analysis was performed on the 7-mer sequence motifs surrounding methylated lysine residues (**Figures S10C-D**). While the amino acid sequence motifs for the Kme sites detected using trypsin/lys-C compared to chymotrypsin digestion are not identical, there is no enrichment for amino acid resides within +/- 3 positions from the central methylated lysine regardless of digestion strategy. Furthermore, there are no pronounced differences in mass, charge, or mass-to-charge ratio of methylated peptides detected following either digestion strategy (**Figure S10E**).

**Figure 4:**
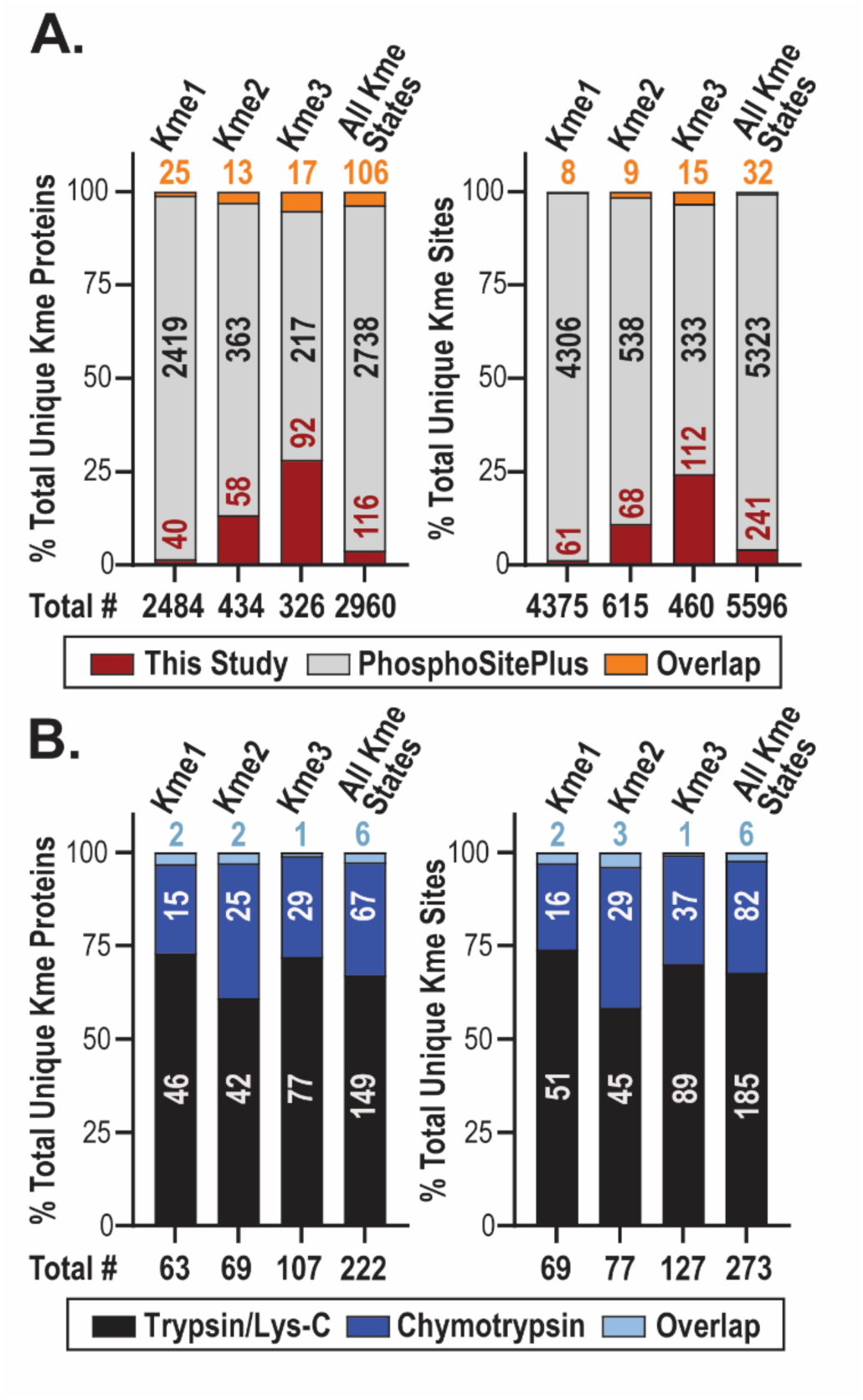
Detection of unique non-histone lysine methylation sites by MS following trypsin/lys-C and chymotrypsin digestion. Number of unique proteins with lysine methylation and unique lysine methylation sites **(A)** detected overall in this study and previously reported in the PhosphoSitePlus® database and **(B)** detected in the trypsin/lys-C and chymotrypsin experiments within this study.

Ultimately, more Kme sites were detected by MS following trypsin/lys-C versus chymotrypsin digestion, and regardless of digestion strategy, most detected peptides with Kme contained 1 additional K/R residue. Potential explanations could be that only a subset of KMTs prefer K- and R-rich sequences, or that the *in vitro* preference of KMTs for K- and R-rich sequences is an artifact and does not translate to substrates methylated in cells. It is also possible that the K- and R-rich sequences are lost in other steps of sample preparation or not retained during the chromatography steps. Nonetheless, the use of both digestion strategies resulted in a more comprehensive detection of Kme sites by MS, as novel sites were identified using both approaches.

### Elucidation of unique non-histone lysine methylome signatures across differentiation stages

Of the 303 methylated peptides corresponding to 273 unique Kme sites detected in this study, 146 methylated peptides corresponding to 127 unique Kme sites (103 unique to trypsin/lys-C MS and 19 unique to chymotrypsin MS) on 98 unique proteins were quantified and normalized to the abundance of the corresponding protein to account for changes in protein abundance across differentiation (**Figures S11 and S12A**). Of the quantified methylated peptides, 79 methylated peptides corresponding to 74 unique Kme sites (70 unique to trypsin/lys-C MS and 4 unique to chymotrypsin MS) on 57 unique proteins are differentially abundant across differentiation (ANOVA padj<0.05, Tukey pairwise padj<0.05) (**Figure 5A and Table S1**).

**Figure 5:**
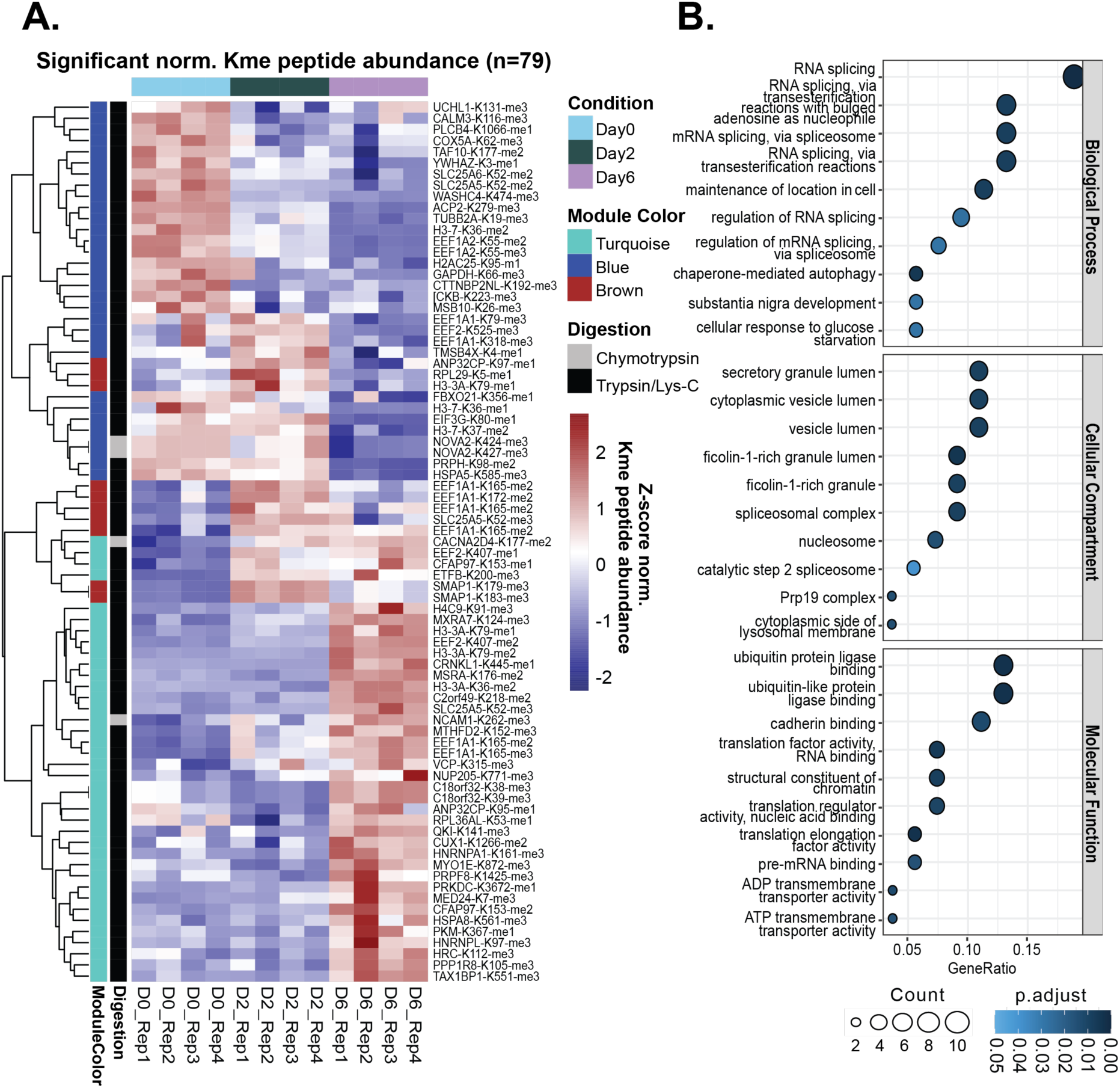
Elucidation of unique non-histone lysine methylome signatures across differentiation stages. **(A)** Heatmap of differentially abundant (ANOVA padj<0.05 followed by Tukey pairwise padj<0.05) Kme peptides (n=79) corresponding to 74 unique Kme sites quantified across LUHMES differentiation. Colors represent the z-score of normalized Kme peptide abundance. Rows (Kme sites) and columns (differentiation sample) are clustered by Euclidian distance. Represented to the left of the rows is the weighted gene correlation network analysis (WGCNA) co-expression color module associated with a given lysine methylated peptide, as well as the digestion strategy following which the Kme peptide was quantified. **(B)** Enriched gene ontology (GO) terms for 57 unique proteins corresponding to differentially abundant Kme sites (p<0.05).

GO term enrichment analysis of all 98 unique proteins corresponding to the 127 quantified Kme sites reveals an enrichment for biological processes mainly related to RNA splicing, molecular functions including ubiquitin and cadherin binding, and cellular compartments including spliceosome complex and ribosome (**Figure S12B**). GO term enrichment analysis of all 57 unique proteins corresponding to the 74 differentially abundant Kme sites reveals an enrichment for biological processes like those identified for all quantified sites, highlighted by RNA splicing (**Figure 5B**). However, proteins associated with differentially abundant Kme sites are also enriched for processes related to dopaminergic and neural development (substantia nigra/neural nucleus development). Interestingly, the substantia nigra is a midbrain nucleus that has dopaminergic projections to the basal ganglia, which modulate functions including cognition and learning, reward, and movement. Damage to the subcortical nuclei of the basal ganglia can result in neurodegenerative and neurodevelopmental disorders^43^.

WGCNA analysis of the 74 differentially abundant Kme sites reveals 3 distinct co-expression modules of methylated peptides with enriched expression on differentiation days 0, 2, or 6 (**Figures S13A-B**). Proteins corresponding to methylated peptides with enriched expression on differentiation day 0 (blue module, n=26 proteins corresponding to 31 Kme peptides) have an enrichment for the biological processes of substantia nigra/neural nucleus development, transmembrane transport, and maintenance of location in cell, and molecular functions associated with ubiquitin protein ligase binding, translational regulation (RNA and nucleic acid binding), and actin monomer binding (**Figure S13C**). There were no enriched processes identified for proteins corresponding to methylated peptides with enriched expression on differentiation day 2 (brown module, n=6 proteins corresponding to 10 Kme peptides). Proteins corresponding to methylated peptides with enriched expression on differentiation day 6 (turquoise module, n= 32 proteins corresponding to 38 Kme peptides) have an enrichment for cellular compartments including the lumen of vesicles and secretory granules (**Figure S13C**). Enrichment of these cellular compartments could suggest a role in transport, storage, or secretion, as secretory granules store biologically active secretory products - including neuropeptides, growth factors, and hormones - for release upon stimulation in endocrine and neuroendocrine cells^44^.

Overall, quantified methylation events occur on diverse non-histone proteins (**Table S2**). Notable examples are discussed below, including methylation of signaling proteins (Calmodulin-3), cytoskeletal proteins (Beta tubulin and Cofilin-1), RNA splicing factors (Nova-2 and Crooked neck-like protein 1), and transcription factors (Homeobox protein cut-like 1).

Calmodulin-3 is among the proteins (n=3) corresponding to methylated peptides associated with the GO terms “substantia nigra/neural nucleus development.” Two tryptic/lys-C peptides were quantified for tri-methylated calmodulin-3 at K116 (CaM3-K116-me3), one with a statistical decrease in abundance on differentiation days 2 and 6 relative to day 0 (**Figure 6A**), and one with the same trend but not differentially abundant. Calmodulin is a calcium (Ca2+) sensor that mediates calcium-dependent signaling through interaction with diverse substrates. Calmodulin regulates most of its binding partners in a calcium-independent manner, whereby Ca2+ binding to calmodulin induces conformational changes that expose hydrophobic patches for substrate binding^45^. One of the major targets of calmodulin is CamKII (Calcium/calmodulin-dependent serine/threonine protein kinase II). Upon Ca2+ binding to calmodulin, calmodulin undergoes a conformational change that confers its binding to CamKII, triggering CamKII autophosphorylation and activation^46^. Through its phosphorylation activity on a wide range of substrates in the brain, CamKII regulates diverse neuronal activities, including: neurotransmitter synthesis and release, modulation of ion channel activity, long-term potentiation, gene expression, learning and memory, and synaptic plasticity^47^. CaM3-K116-me3 (at times referenced in the literature as K115 based on previous annotation), mediated by Calmodulin-lysine N-methyltransferase (CAMKMT), is a conserved modification across a variety of species. Tri-methylation of calmodulin at K116, which resides within the third of four EF-hand calcium binding domains, has been demonstrated to attenuate its activation of calmodulin-dependent NAD kinase *in vitro*^48^. Based on the literature, calmodulin-3 K116 tri-methylation in LUHMES cells disrupts calmodulin binding to and activation of its targets, potentially impacting downstream calcium-mediated signaling pathways.

**Figure 6:**
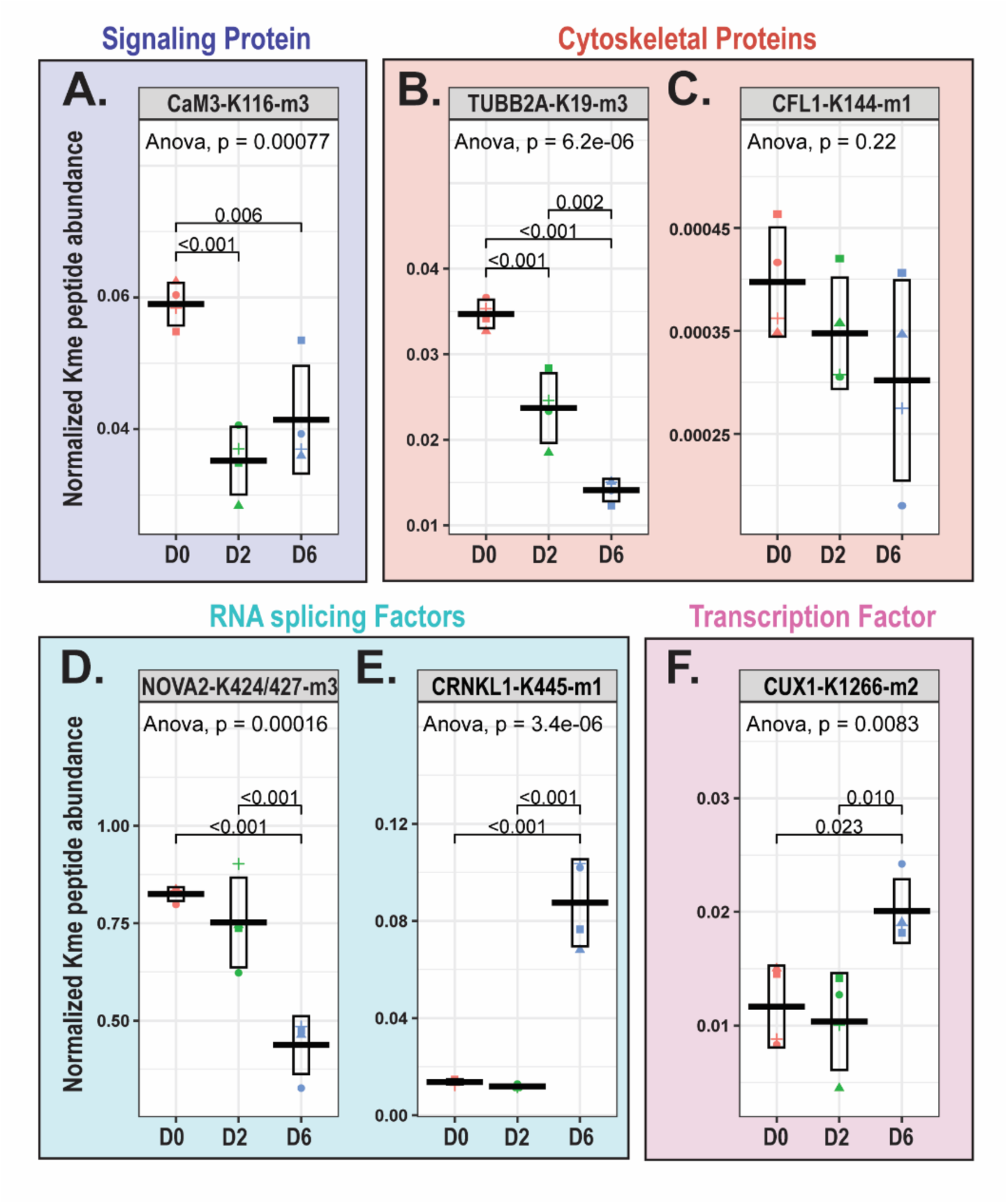
Lysine methylation of diverse non-histone proteins. Boxplots displaying mean+/-standard deviation of the normalized Kme peptide abundance for **(A)** tri-methylated Calmodulin-3 at K116 (CaM3-K116-m3), **(B)** tri-methylated β-tubulin-2A at K19 (TUBB2A-K19-m3), **(C)** mono-methylated Cofilin-1 at K144 (CFL1-K144-m1), **(D)** tri-methylated Nova-2 at K4224/427(NOVA2-K424/427-m3), **(E)** mono-methylated Crooked-neck like protein 1 at K445 (CRNKL1-K445-m1), and **(F)** di-methylated Homeobox protein cut-like 1 at K1266 (CUX1-K1266-m2). ANOVA followed by Tukey pairwise comparison. Each data point shape represents a different biological replicate.

Tubulin proteins are the major constituents of microtubule polymers, which are essential for neuronal structure and polarity, cell division, and intracellular transport^49^. Across LUHMES differentiation, there is decreased abundance of tri-methylated β-tubulin-2A at K19 (TUBB2A-K19-me3) (**Figure 6B**), which resides within a GTP-binding domain conserved among tubulin proteins^50^. Tubulin GTP hydrolysis activity is critical for regulating microtubule dynamics^50^, and interestingly, lysine methylation within GTP-binding domains has been demonstrated to alter GTPase activity^51^. Functional annotation of β-tubulin methylation has not been reported (though K19 methylation has been reported in PhosphoSitePlus®); nonetheless, there are reports of functional impacts of α-tubulin methylation. For instance, SETD2-mediated tri-methylation of α-tubulin at K40 (within its GTP-binding domain) promotes microtubule formation, through tubulin nucleation and microtubule polymerization, and is required for cortical neuronal polarization and migration^13^. Thus, it is possible that methylation at K19, within the GTP-binding domain of β-tubulin-2A, could impact GTP binding/hydrolysis and microtubule polymerization. Coffilin-1 (CFL1) is another cytoskeletal protein methylated in LUHMES, but it does not change in abundance (**Figure 6C**). CFL1 is methylated at lysine 144 (CFL1-K144-me1), which resides within its actin-depolymerizing factor (ADF) homology domain. Cofilin-1 binds to F-actin and mediates its depolymerization^52^; this activity has been reported to regulate growth cone actin dynamics and dendritic spine morphology^53,54^. The activity of cofilin-1 is highly regulated by its phosphorylation and acetylation status. Cofilin-1 depolymerization activity is inhibited by its phosphorylation at serine 3, and inhibition of cofilin-1 phosphorylation results in actin depolymerization and the promotion of neurite outgrowth^55,56^. Cofilin-1 acetylation and phosphorylation appear to be mutually exclusive, where acetylation of cofilin-1 (at K33, K44, K96 and/or K144 residues) may prevent its serine 3 phosphorylation^57^. The phosphorylation/acetylation status of cofilin-1 impacts its interaction with actin-binding protein cortactin, with which it regulates microtubule dynamics; acetylation of cofilin-1 enables its interaction with cortactin while phosphorylation prevents it^57^. Similar to the reported impacts of acetylation of cofilin-1 at K144, methylation at this residue may impact cofilin-1 depolymerization activity and/or substrate interaction.

Several proteins implicated in RNA splicing were differentially methylated across differentiation. Tri-methylation of RNA-binding protein Nova-2 at lysine residues 424 and 447 (NOVA2-K424/427-me3) decreased on differentiation day 6 relative to days 0 and 2 (**Figure 6D**). Nova proteins are neuronal-specific alternative splicing factors, and Nova-2 has been found to regulate alternative splicing events of genes involved in axon guidance, synaptic formation, and synaptic plasticity^58,59^. Nova-2 contains three RNA binding (K homology, KH) domains, and the Nova-2 methylation events quantified in this study (tri-methylation of K424 and K427) occur within its KH 3 domain. Methylation at these sites could impact Nova 2 RNA binding, interaction with other splicing regulators, and alternative splicing. Notably, Nova-2 K424/427-me3 was one of four differentially abundant Kme sites quantified following chymotrypsin digestion, but not detected in the trypsin/lys-C MS experiment. Manual inspection of the sequence context surrounding this site shows it is unlikely to be detected by trypsin/lys-C digestion. Tri-methylation of both lysine residues increases the size and bulkiness of the side chain, and steric hindrance would likely prevent trypsin/lys-C cleavage after the modified sites^60^. Even if trypsin/lys-C cleavage were to occur, the predicted tryptic peptide would likely be too long for detection by MS, as there is not another lysine or arginine residue within proximity to the methylated sites. On differentiation day 6 relative to days 0 and 2, there is an increased abundance of mono-methylated Crooked-neck like protein 1 at K445 (CRNKL1-K445-me1) (**Figure 6E**). Crooked neck 1, a pre-MRNA splicing factor, is required for glial cell migration and axonal wrapping in *Drosophila* and, through interaction with RNA-binding protein Held out wings (HOW), controls differentiation through facilitation of splicing of genes involved in cellular junction assembly^61^. Crooked neck 1 is methylated at lysine residue 445, which resides within repeat region 7 of its 17 tandem HAT (Half-a-Tetratricopeptide) repeat regions. Evidence suggests that HAT repeat regions can bind RNA and that this binding impacts regulation of gene expression^62^. HAT repeat regions have also been demonstrated to bind peptides^63,64^. It has been reported that the direct interaction between the HAT repeat domain of small subunit processing component Utp6 and a peptide within RNA-associated protein Utp21 is essential for pre-rRNA processing and cell growth^63^. It was proposed that disruption of the Utp6-Utp21 interaction reduces the efficiency of small subunit processome assembly and leads to downstream inhibition of ribosome biogenesis. The HAT 7 repeat region, containing K445, of crooked neck 1 is predicted to mediate interaction with heat shock protein HSP90 (Uniprot manual assertion inferred from sequence similarity)^24^; it is possible that K445 methylation could impact interaction of crooked neck 1 with proteins, including HSP90, or with RNA, which could impact its downstream RNA splicing activity.

Methylation of transcription factors also changed during differentiation. For example, di-methylation of K1266 on CUX1 (CUX1-K1266-me2) increased on D6 (**Figure 6F**). CUX1 is a member of the homeodomain transcription factor family of proteins that bind distinct DNA sequences. CUX1 regulates transcription of genes required for cell migration and invasion, and acts as a transcriptional repressor or activator depending on its interaction partners^65^. CUX1, along with CUX2, promotes dendritic branching and spine development in upper-layer cortical neurons in part through transcriptional repression of Xlr chromatin remodeling genes through direct DNA binding^66^. Methylation of CUX1 at K1266, which resides within its homeodomain, could impact its DNA binding activity and downstream transcriptional regulation.

### Methylation of lysine residues with reported neurodevelopmental disorder-associated clinical variants

To determine if lysine methylation events quantified in this study occur at lysine residues that are of previously reported clinical relevance, we probed the ClinVar database^25^. Interestingly, we found clinical variants, associated with neurodevelopmental disorders, at a subset of lysine residues methylated in this study, including K462 of Heterogeneous nuclear ribonucleoprotein U (HNRNPU-K462), K200 of Electron transfer flavoprotein subunit beta (ETFB-K200), and K55 of Elongation factor 1-alpha 2 (EEF1A2-K55) (**Table S3**).

#### HNRNPU-K462

Heterogeneous nuclear ribonucleoprotein U (HNRNPU) is a DNA- and RNA-binding protein with reported functions in chromatin organization, alternative splicing, transcription regulation, X-linked transcriptional silencing during X-inactivation, and genomic stability^67^. Knockout of HNRNPU is embryonic lethal in mice and appears to be essential for cortical development^68^. Conditional truncation of HNRNPU in the developing cortex results in reduced cortex volume, death of neural progenitor cells and postmitotic neurons, and dysregulated expression and alternative splicing of genes critical for brain development and function, including those involved in cell survival and motility, cytoskeleton organization, and synapse formation^68^.

According to the Simons Foundation Autism Research Initiative (SFARI) database^6^ *, HNRNPU* is determined to be a high-confidence syndromic gene (category 1, Syndromic; 1S) associated with autism spectrum disorder (ASD). *HNRNPU* variants are largely associated with phenotypes including global developmental delay, moderate to severe intellectual disability, early onset seizures, and dysmorphic features^69^. Of note, there is a reported ClinVar missense variant for *HNRNPU* at lysine residue 462 (p.Lys462Arg), the residue tri-methylated in our study. This variant is of uncertain significance for Developmental and epileptic encephalopathy, 54 (ClinVar Accession: VCV001943455.4).

In our study, tri-methylated HNRNPU at K462 does not change in abundance across differentiation (**Figure 7A**). The methylated residue, K462, resides within the B30.2/SPRY domain, which mediates protein-protein interactions^70^. HNRNPU is reported to have several ubiquitination sites^22^, including: K352 and K464, which fall in the B30.2/SPRY domain, and K524 and K565, which fall in the ATPase domain. Ubiquitination of HNRNPU, mediated by cell cycle protein CDC20, is required for its interaction with chromatin-organizer protein CTCF^71^. Lysine to arginine mutation of HNRNPU residues 352/464/524/565 results in decreased HNRNPU ubiquitination and reduced interaction between HNRNPU and CDC20, as well as HNRNPU and CTCF^71^. It is possible that tri-methylation of K462 impacts ubiquitination on the proximal K464, or vice versa, through post-translational modification crosstalk, and that this methylation event could impact binding of HNRNPU to interaction partners.

**Figure 7:**
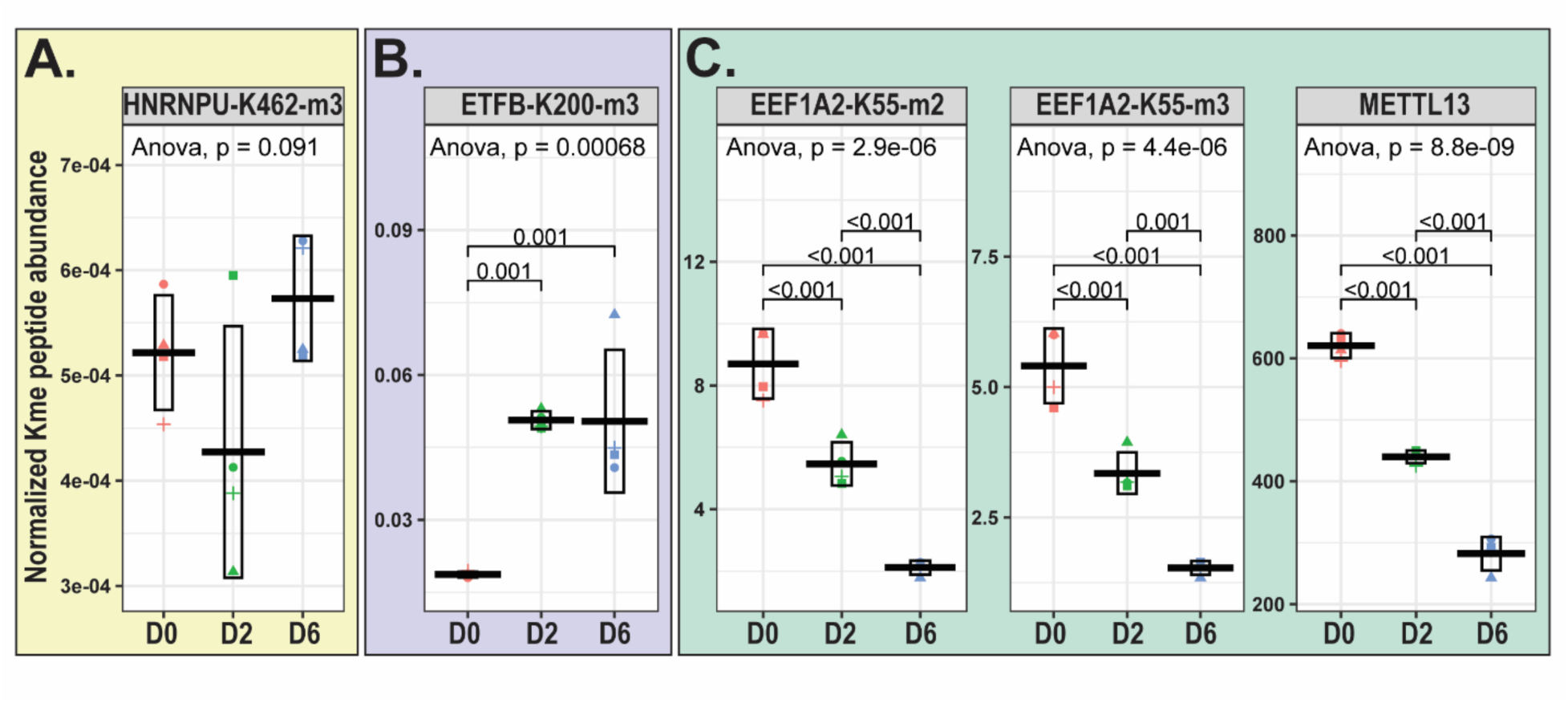
Methylation of lysine residues with reported neurodevelopmental disorder-associated clinical variants. Boxplots displaying mean+/-standard deviation of the normalized Kme peptide abundance for **(A)** tri-methylated Heterogeneous nuclear ribonucleoprotein U at K462 (HNRNPU-K462-m3), **(B)** tri-methylated Electron transfer flavoprotein subunit beta at K200 (ETFB-K200-m3), and **(C)** di- and tri-methylated Elongation factor 1-alpha 2 at K55 (EEF1A2-K55-m2/m3), along with protein abundance of the corresponding lysine methyltransferase METTL13. ANOVA followed by Tukey pairwise comparison. Each data point shape represents a different biological replicate.

#### ETFB-K200

Electron transfer flavoprotein subunit beta (ETFB) is part of the heterodimeric ETF protein complex that functions in maintaining cellular redox balance. It facilitates electron transfer from various metabolic enzymes, including acyl-CoA dehydrogenases, and mitochondrial processes, such as fatty acid oxidation and amino acid catabolism, to the respiratory chain, ensuring proper oxidative phosphorylation and cellular respiration^72^. A metabolic shift toward energy metabolism, including oxidative phosphorylation and mitochondrial function and biogenesis, has been reported to be associated with neuronal differentiation progression and cell fate determination^73–75^.

According to the SFARI database, *ETFB* is a strong candidate (category 2) ASD gene. In one study, *ETFB* was identified in an ASD whole-exome sequencing study as a gene that is enriched in variants that are likely to affect the risk of ASD^76^. ETFB clinical variants have also been associated with multiple acyl-CoA dehydrogenation deficiency, which clinically presents with neonatal onset with metabolic decompensation with/without congenital anomalies, or later onset with milder presentations^77^. There are several reported ClinVar variants for ETFB at K200, the residue tri-methylated in our study, associated with Multiple acyl-CoA dehydrogenase deficiency. Among them is a one-base pair deletion splice acceptor variant [NM_001985.3(ETFB):c.598-1del]. There are two ClinVar classifications of pathogenicity for this variant-one classification is of uncertain significance, while the other is likely pathogenic for Multiple acyl-CoA dehydrogenase deficiency (ClinVar Accession: VCV000808634.30). There is also a reported ClinVar missense variant for ETFB at K200 (p.Lys200Glu) of uncertain significance for Multiple acyl-CoA dehydrogenase deficiency (ClinVar Accession: VCV000459960.8).

In this study, tri-methylation of ETFB at K200 (ETFB-K200-me3) increases in abundance across LUHMES differentiation (**Figure 7B**). ETFB K200 resides within its mitochondrial dehydrogenase recognition loop critical for mediating interactions with proteins and co-factors in the electron transport chain^72^. METTL20-mediated methylation of ETFB K200, as well as K203, has been demonstrated to inhibit ETFB-mediated electron transfer from acyl-CoA dehydrogenases^78^. Thus, it is rational that ETFB-K200 methylation in LUHMES cells could affect ETFB-mediated electron transfer, influencing mitochondrial function and energy production which ultimately impact neuronal differentiation.

#### EEF1A2-K55

Elongation factor 1-alpha (EEF1A), one of the most abundant proteins in eukaryotic cells, canonically functions in translation elongation by delivering amino-acylated transfer RNAs (aa-tRNAs) to the elongating ribosome in a GTP-dependent manner during protein biosynthesis^79^. It has also been reported to participate in many other processes, including cell migration, signal transduction, nucleocytoplasmic trafficking, and cytoskeletal organization and remodeling^80^. Mammalian EEF1A has two isoforms-EEF1A, which is expressed ubiquitously, and EEF1A2, which has enriched expression in muscle and neurons^81^. Interestingly, the activities of the EEF1A2 isoform specifically appear to be critical in neurons. Mohamed et al. postulate that EEF1A2 activities coordinate the intersection of translation and the actin cytoskeleton^82^. This group found that the introduction of three common autism- and epilepsy-associated variants of *EEF1A2* (G70S, E122K, and D252H) results in decreased de novo protein synthesis and reduced translation elongation in HEK cells. Introduction of these same mutations in primary cortical neurons also results in decreased de novo protein synthesis and altered neuronal morphology. The authors posit that the altered neuronal morphology upon *EEF1A2* mutation is due to increased EEF1A2 tRNA binding capacity (which could result in broad sequestration of tRNAs, decreased tRNA availability, and reduced rate of translation elongation) and decreased actin-bundling activity (which could disrupt the connection between the actin cytoskeleton and mRNA translation). Additionally, EEF1A2 appears to be essential *in vivo*. EEF1A2 was found to be essential for post-weaning survival in the mouse model known as “wasted,” which is characterized by deletion of *EEF1A2*, and loss of EEF1A2 function is attributed to the phenotypes, including motor neuron degeneration, of wasted mice^83^.

According to the SFARI database, *EEF1A2* is considered to be a syndromic gene (category S) for ASD. In several reports, variants in *EEF1A2* have been associated with autistic features, intellectual disability, and epilepsy^84–86^. Interestingly, there is a reported ClinVar missense variant for *EEF1A2* at lysine residue 55 (p.Lys55Arg), the residue di- and tri-methylated in our study. This variant is of uncertain significance for Neurodevelopmental disorder (ClinVar Accession: VCV001701882.1).

In our study, we quantify both di-methylation and tri-methylation of EEF1A2 at K55 (EEF1A2-K55-me2 and EEF1A-K55-me3), and the abundance of both methyl states decreases across LUHMES differentiation (**Figure 7C**). Furthermore, in this study, we were able to quantify the abundance of METTL13, the KMT that mediates EEF1A2 methylation^51^, and, as expected, METTL13 abundance also decreases across LUHMES differentiation (**Figure 7C**). METTL13-mediated EEF1A methylation at K55, which resides within its translational (tr)-type GTP-binding domain, has been reported to increase EEF1A GTPase activity *in vitro* and stimulate protein synthesis in cancer cells^51^. Based on this reported impact of EEF1A K55 methylation, as well as the reported role of EEF1A2 in protein synthesis in neurons^82^, it is rational to hypothesize that METTL13-mediated EEF1A2 methylation at K55 could impact its GTPase activity and downstream regulation of translation elongation and protein synthesis in LUHMES cells.

## Discussion

Lysine methylation of histone proteins has been reported to regulate neuronal differentiation, primarily by regulating gene expression. Moreover, quantitative profiling of histone modifications across neuronal differentiation has revealed differentiation-stage-specific expression patterns of histone lysine methylation^12^. In contrast, there are limited reports on the roles of non-histone lysine methylation in neuronal differentiation. In this work, we simultaneously profiled global protein abundance and lysine methylation sites across LUHMES differentiation days 0 (neural progenitor stage), 2 (early differentiation stage), and 6 (post-mitotic, dopaminergic-like neuron) using Tandem Mass Tag (TMT) mass spectrometry (MS) without immunoaffinity enrichment for lysine methylation. To our knowledge, this is the first report of quantitative profiling of global non-histone lysine methylation across neuronal differentiation.

Overall, we quantified 5,568 unique proteins, most of which showed differentiation-stage-specific expression. Consistent with our previous work profiling global lysine methylation using TMT-based MS workflows^40^, we detected 273 unique Kme sites on 222 unique proteins, of which ∼90% are novel. Of the 273 Kme sites detected, 127 unique Kme sites on 98 unique proteins were quantified and normalized to protein abundance. Among these, 74 sites are differentially abundant across differentiation, revealing unique non-histone lysine methylome signatures across differentiation stages. We identified lysine methylation events on a range of diverse non-histone proteins with reported roles in neuronal differentiation and neurodevelopment. Among the methylated proteins are signaling proteins (Calmodulin-3), cytoskeletal proteins (Beta tubulin and Cofilin-1), RNA splicing factors (Nova-2 and Crooked neck-like protein 1), and transcription factors (Homeobox protein cut-like 1). Furthermore, we identified methylation events on proteins highly associated with neurodevelopmental disorders (NDDs) at lysine residues for which there are reported NDD-associated clinical variants, including K462 of Heterogeneous nuclear ribonucleoprotein U (HNRNPU-K462), K200 of Electron transfer flavoprotein subunit beta (ETFB-K200), and K55 of Elongation factor 1-alpha 2 (EEF1A2-K55). These clinical variants include missense variants, suggesting that disruption of methylation at these sites could have clinical impacts.

As part of our proteomics approach, we evaluated the detection of Kme sites following two different peptide digestion strategies, trypsin/lys-C or chymotrypsin digestion. We found that ∼68% of all Kme sites detected in this study were uniquely detected following trypsin/lys-C digestion. While more Kme sites were detected following trypsin/lys-C digestion, the complementary use of both digestion strategies ultimately resulted in a more comprehensive detection of Kme sites by MS. Future work will evaluate the use of other strategies to complement trypsin/lys-C digestion for the detection of Kme sites by MS. Overall, this work expands the scope of reported non-histone lysine methylation in a neuronal differentiation model and provides a resource of Kme sites on proteins of biological and clinical interest for future study.

## Data Availability

Data are available via ProteomeXchange with identifier PXD069591.

## Acknowledgments

The research reported in this publication was supported by the National Institute of General Medical Sciences of the National Institutes of Health under award number R35GM147023 (EMC). The content is solely the responsibility of the authors and does not necessarily represent the official views of the National Institutes of Health. The mass spectrometry work was performed by the Indiana University School of Medicine (IUSM) Center for Proteome Analysis (CPA). Acquisition of the IUSM CPA instrumentation used for this project was provided in part by the Indiana University Precision Health Initiative and the IU Simon Comprehensive Cancer Center. The proteomics work was supported, in part, by the Indiana Clinical and Translational Sciences Institute (Award Number UL1TR002529 from the National Institutes of Health, National Center for Advancing Translational Sciences, Clinical and Translational Sciences Award) and, in part, by the IU Simon Comprehensive Cancer Center Support Grant (Award Number P30CA082709 from the National Cancer Institute). The mass spectrometry proteomics data have been deposited to the ProteomeXchange Consortium via the PRIDE^87^ partner repository with the dataset identifier PXD069591 and 10.6019/PXD069591

---

**Figure S1:**
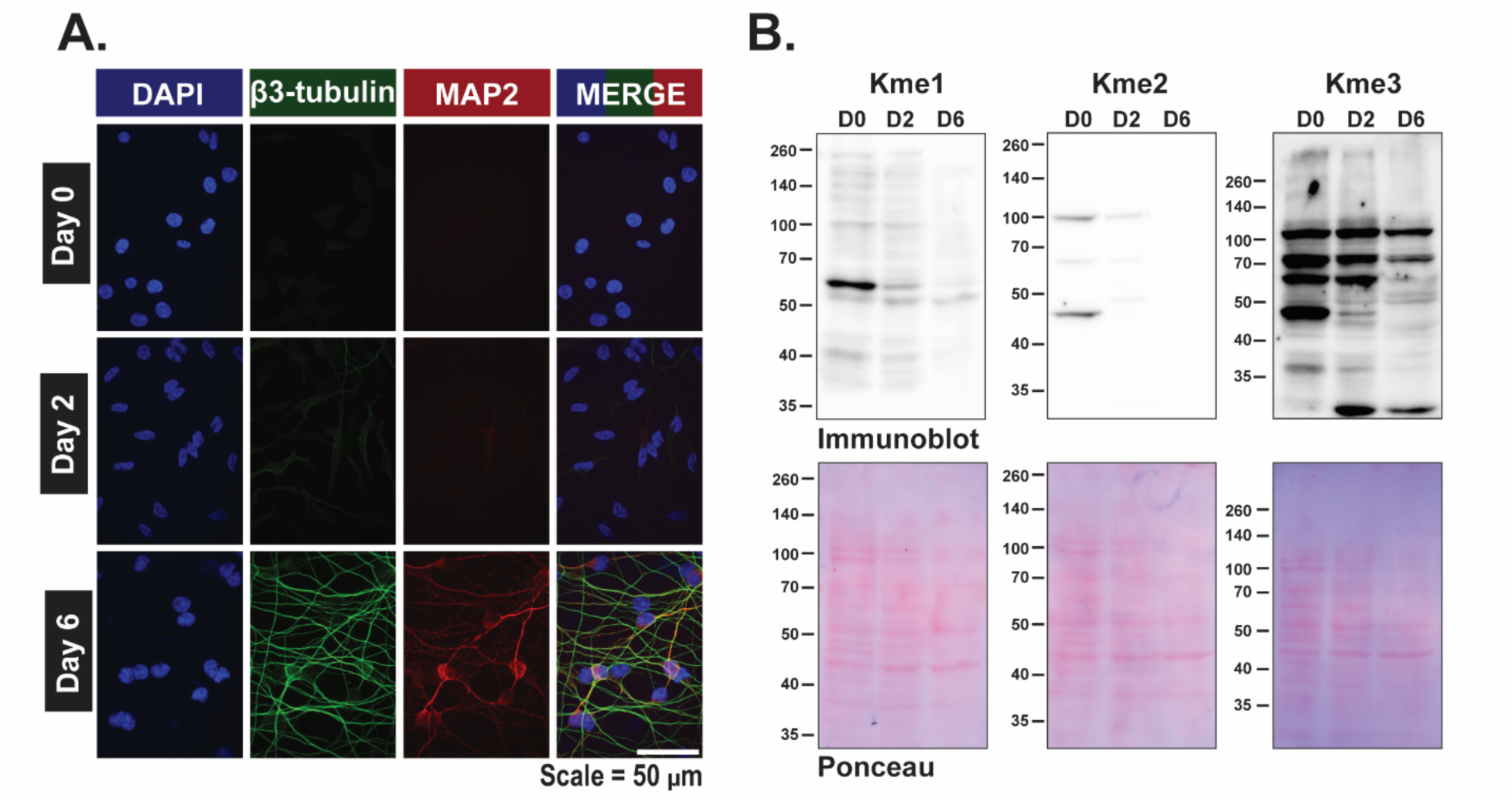
Characterization of LUHMES cell differentiation. **(A)** Immunocytochemistry of undifferentiated, day 2 differentiated, and day 6 differentiated LUHMES cells: β3-tubulin (green), MAP2 (red), and Nucleus (Blue). Scale bar represents 50 µm. **(B)** *In vitro* changes in lysine methylation of non-histone proteins across LUHMES differentiation days 0, 2, and 6 (D0, D2, D6) detected via immunoblotting using pan-lysine methyl antibodies against all three methyl states, mono-(Kme1), di-(Kme2), and tri-(Kme3) methylation. Top panel depicts immunoblotting (25 µg LUHMES lysate loaded per lane) and bottom panel depicts Ponceau Red staining.

**Figure S2:**
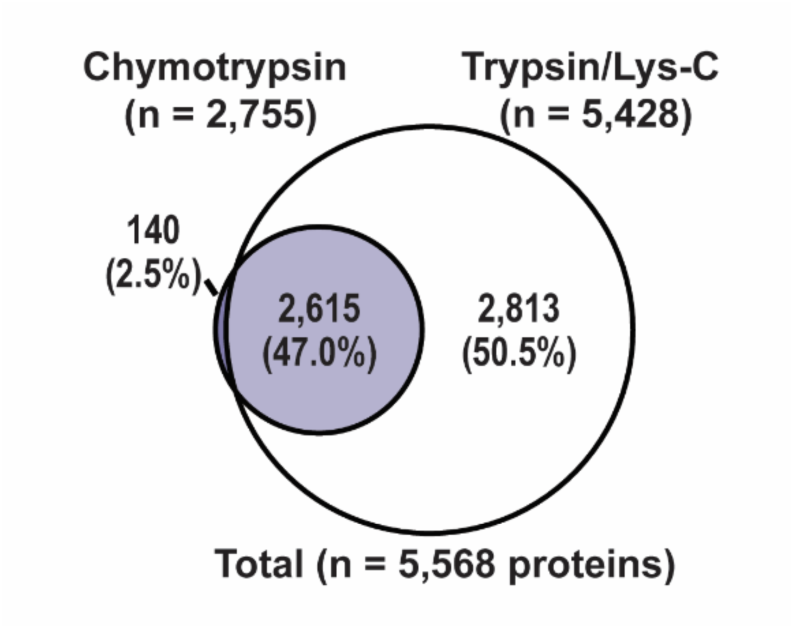
Comparison of the LUHMES proteome from the trypsin/lys-C and chymotrypsin experiments. Venn diagram depicting overlap of proteins quantified in the trypsin/lys-C and chymotrypsin experiments.

**Figure S3:**
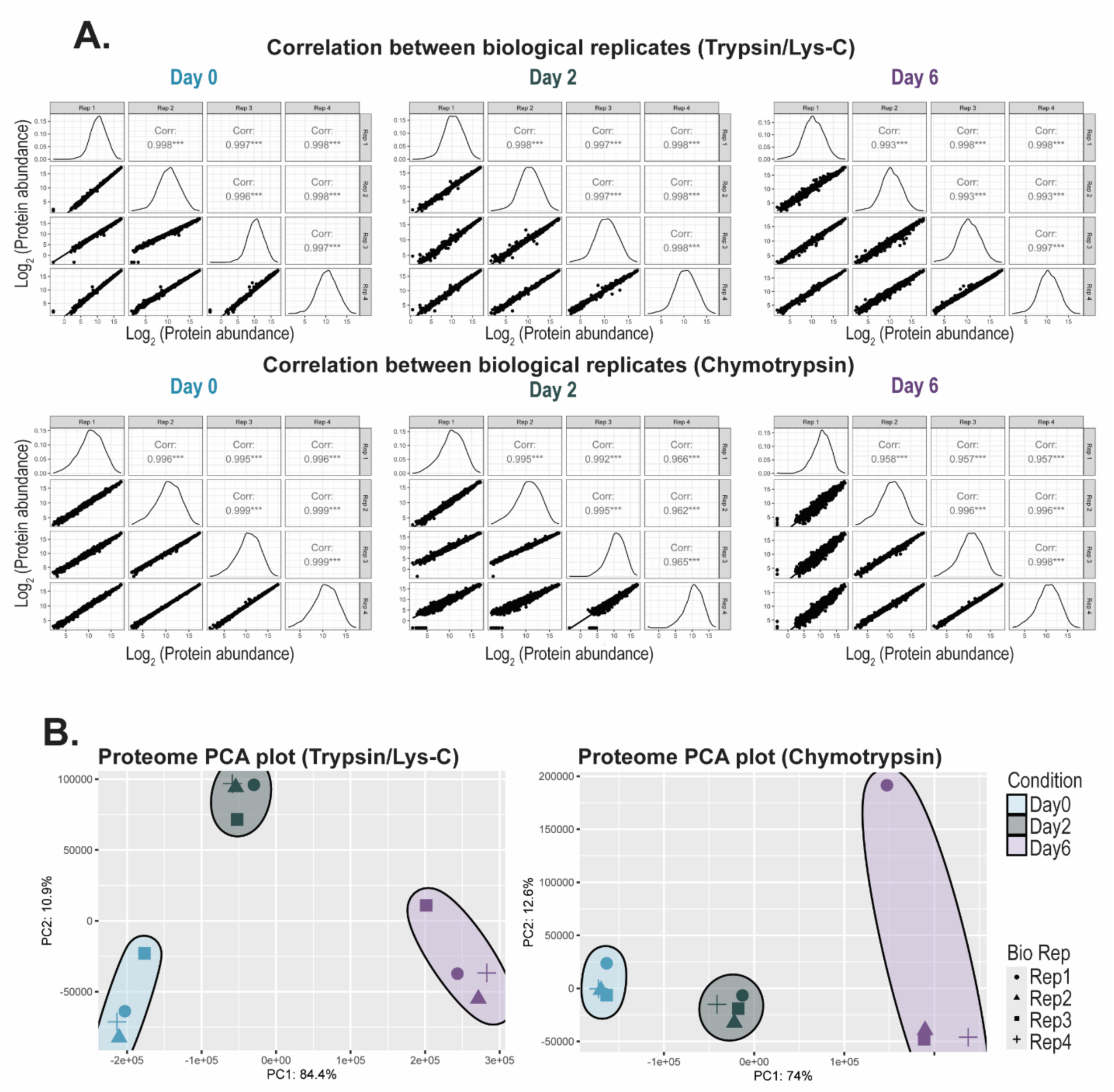
Correlation of protein abundance across biological replicates within the trypsin/lys-C and chymotrypsin experiments. **(A)** Pearson correlation analysis of the average log_2_ abundance of quantified proteins between biological replicates within the trypsin/lys-C experiment (n=5,428) and within the chymotrypsin experiment (n=2,755). **(B)** Principal component analyses (PCA) of the trypsin/lys-C and chymotrypsin proteomes. Each data point shape represents a different biological replicate.

**Figure S4:**
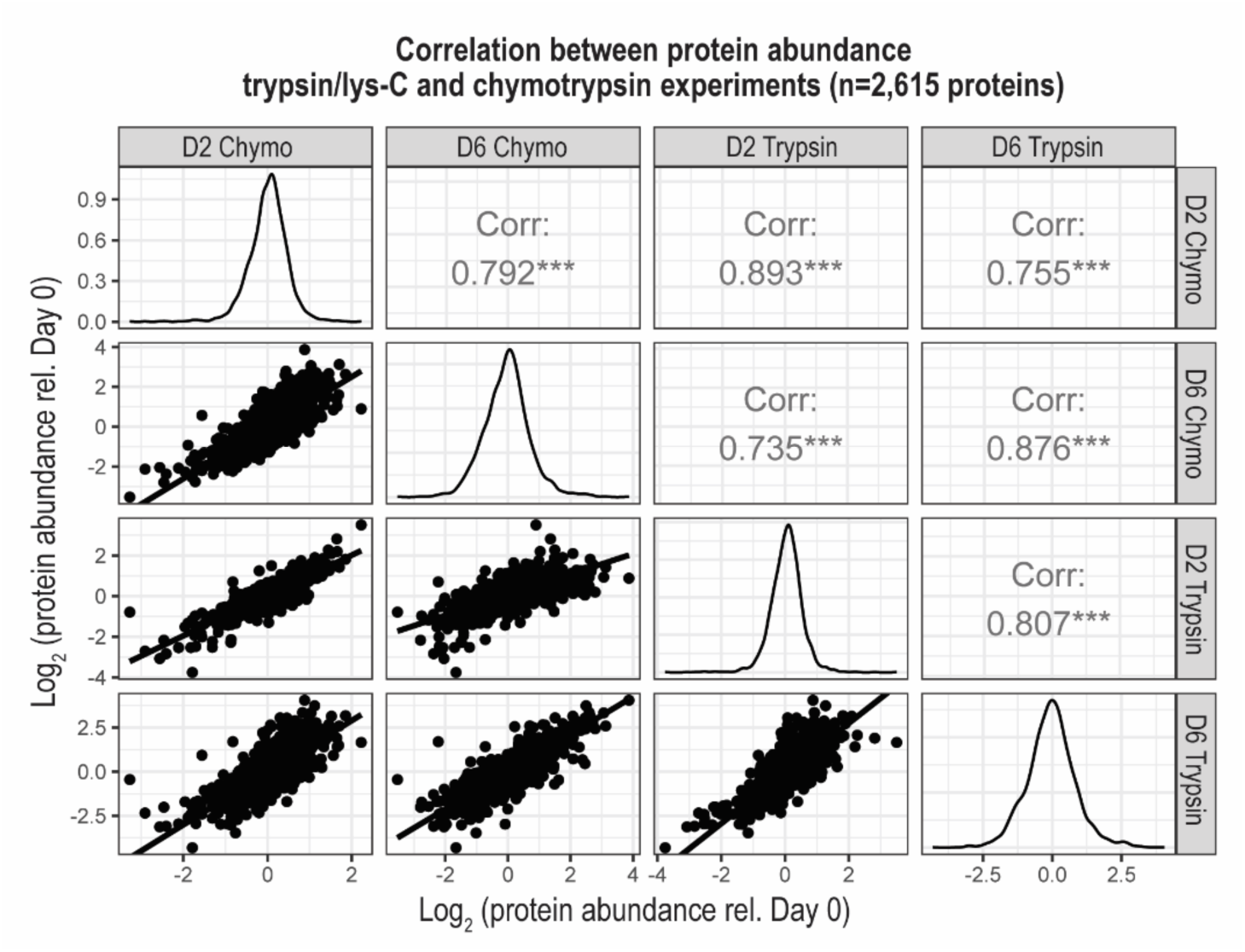
Correlation between abundance of proteins quantified in the trypsin/lys-C and chymotrypsin experiments. Pearson correlation analysis of the average log_2_ protein abundance on differentiation days 2 and 6 relative to day 0 for proteins quantified in both the trypsin/lys-C and chymotrypsin experiments (n=2,615).

**Figure S5:**
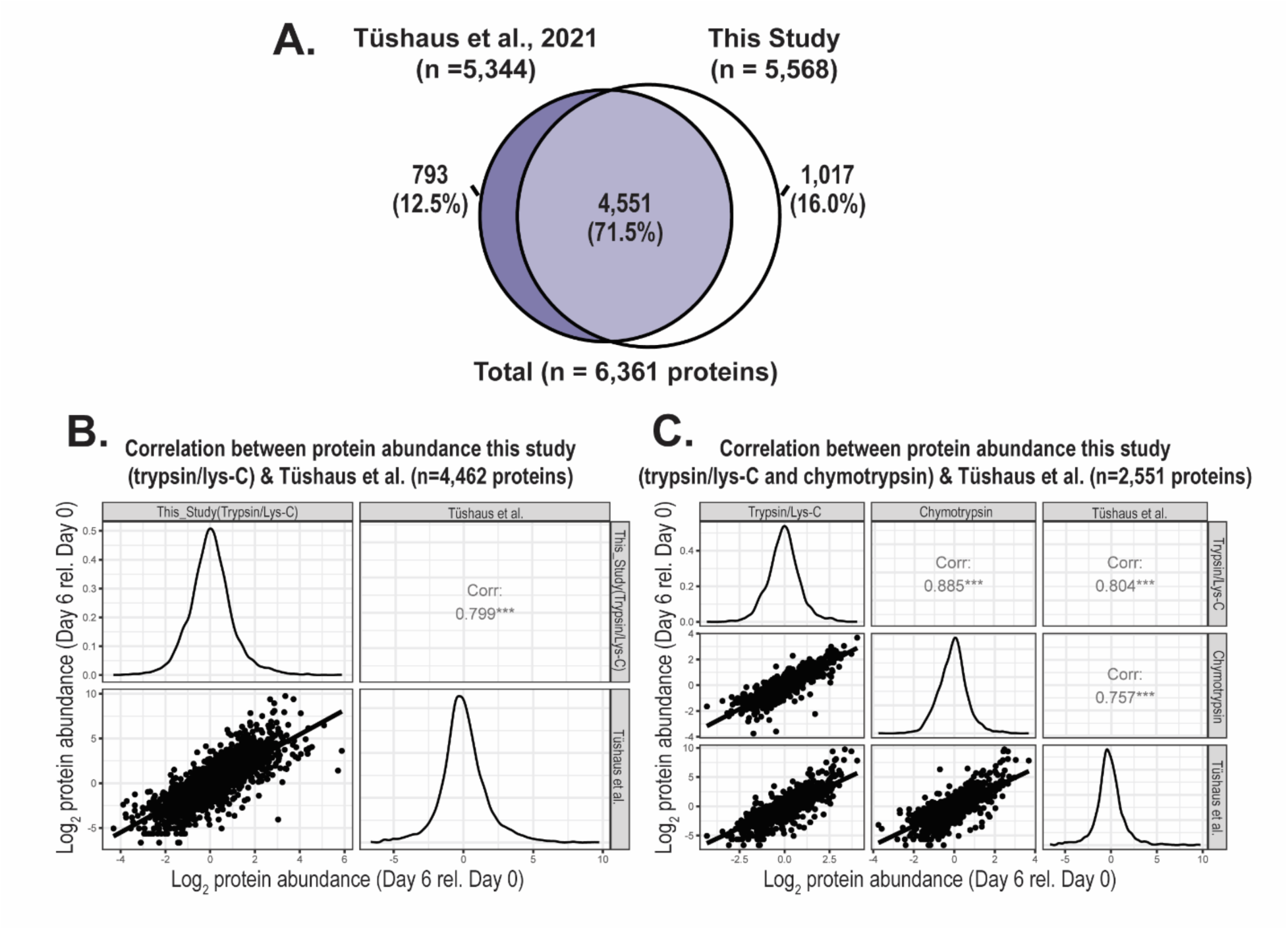
Correlation between the LUHMES proteome from this study and Tüshaus et al. **(A)** Venn diagram depicting overlap of proteins quantified in this study and Tüshaus et al. **(B)** Pearson correlation analysis of the average log_2_ protein abundance on differentiation day 6 relative to day 0 of proteins quantified in Tüshaus et al. and our trypsin/lys-C experiment (n=4,462), and **(C)** proteins quantified in Tüshaus et al. and our trypsin/lys-C and chymotrypsin experiments (n=2,551).

**Figure S6:**
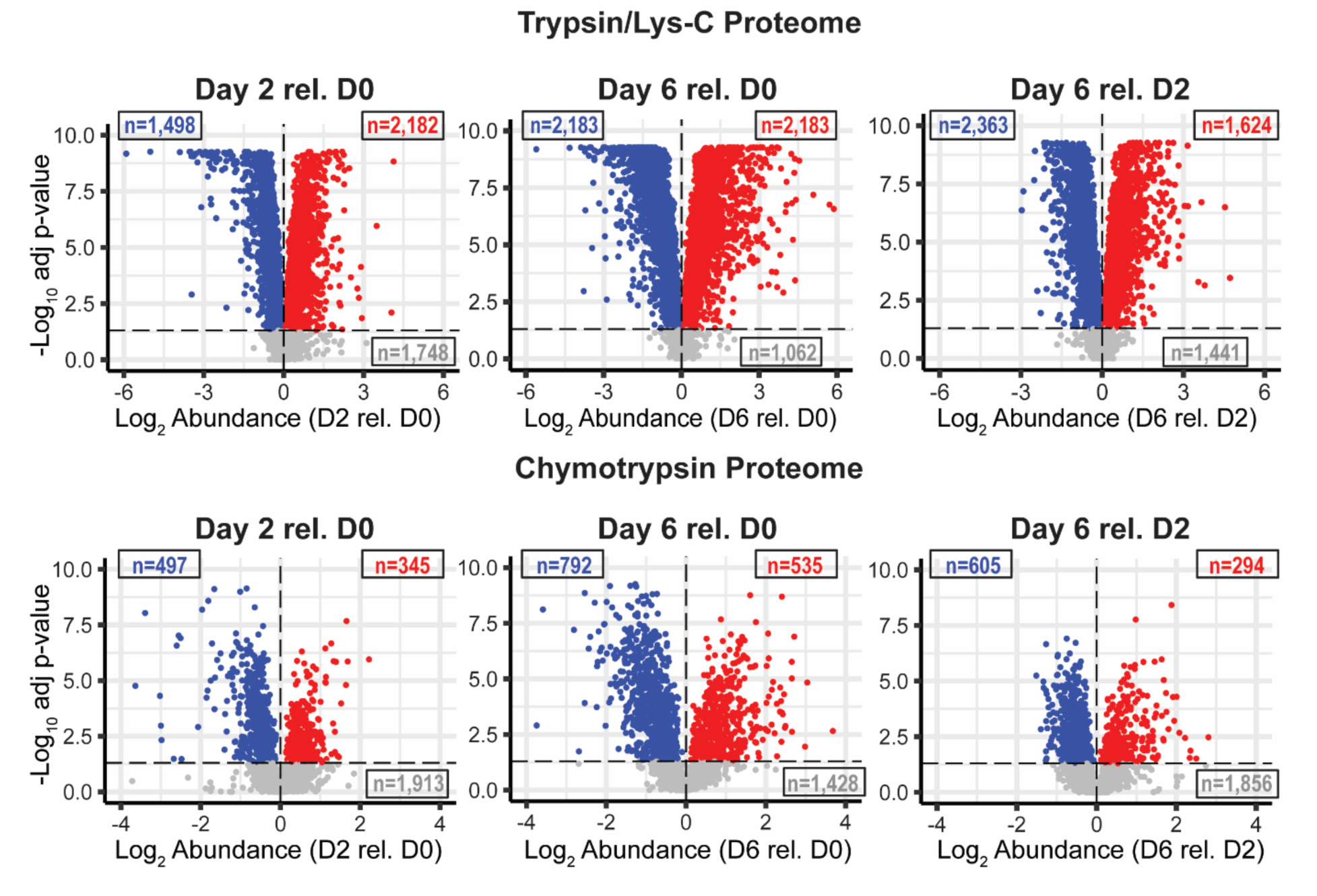
Global protein abundance changes across LUHMES differentiation. Volcano plots displaying log_2_ protein abundance (x-axis) at one day of differentiation relative to another, as depicted. P-values plotted on the y-axis. Significantly (ANOVA padj<0.05 followed by Tukey pairwise padj<0.05) upregulated and downregulated proteins are colored in *red* and *blue*, respectively, while proteins with no statistical change in abundance are colored in *grey*.

**Figure S7:**
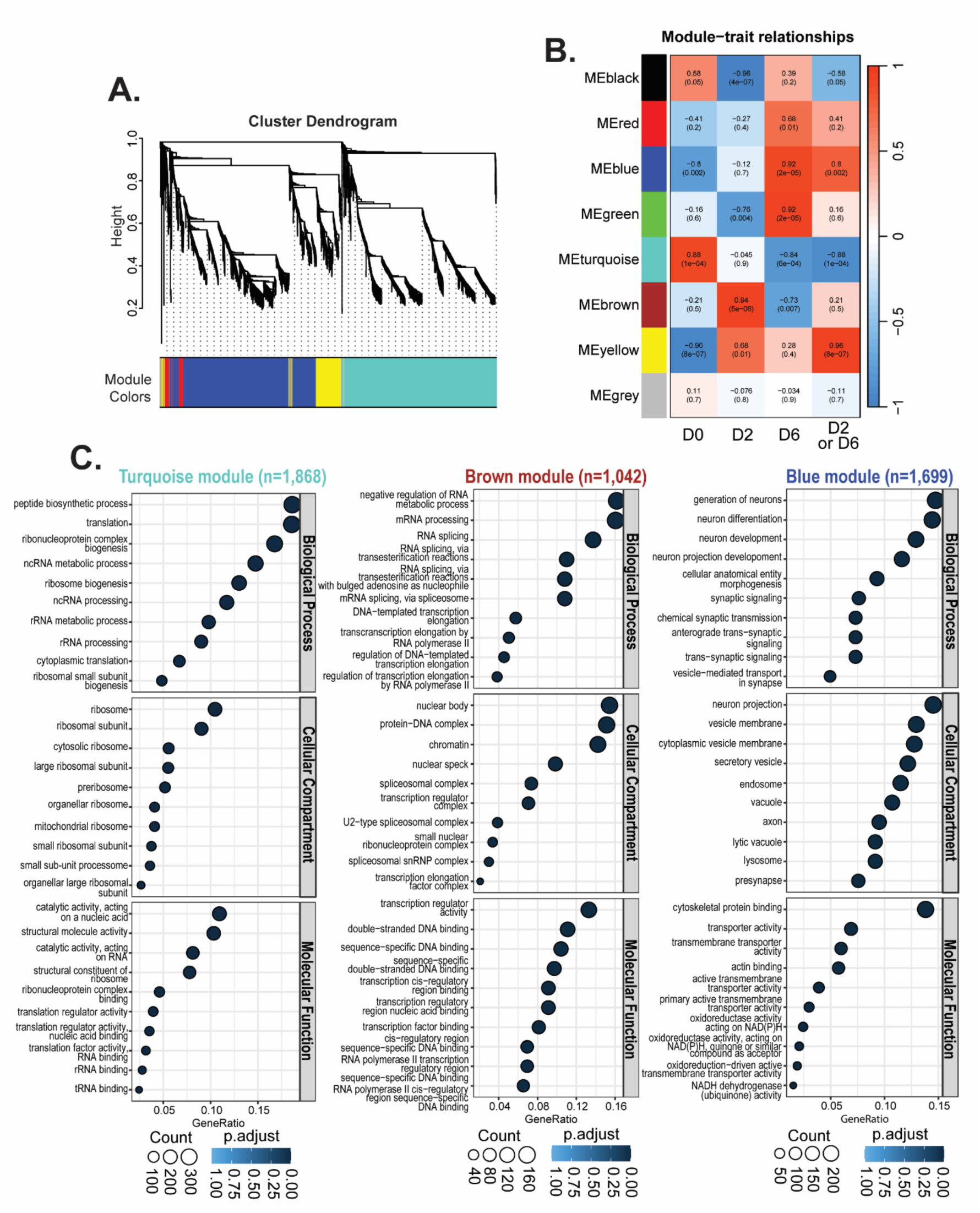
Distinct protein co-expression modules and associated functions revealed through WGCNA and GO terms analyses. **(A)** WGCNA dendrogram and **(B)** module-trait relationship correlation of protein co-expression modules identified by WGCNA analysis of all proteins quantified in the trypsin/lys-C experiment. **(C)** Enriched GO terms for proteins associated with the following modules: turquoise (enriched expression differentiation day 0), brown (enriched expression differentiation day 2), and blue (enriched expression differentiation day 6).

**Figure S8:**
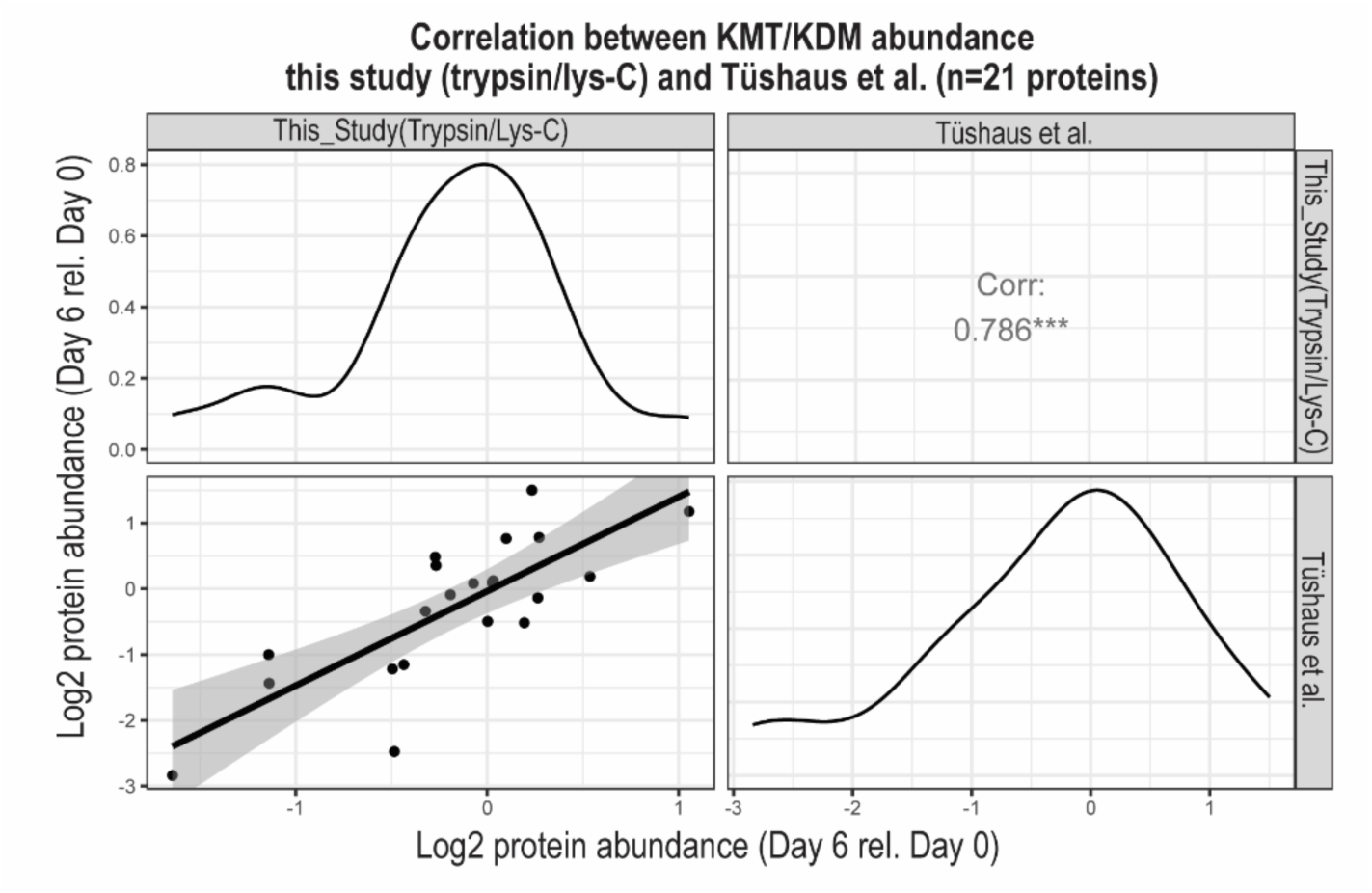
Correlation between abundance of KMTs/KDMs quantified in this study and Tüshaus et al. Pearson correlation analysis of the average log_2_ abundance of KMTs/KDMs quantified in both the trypsin/lys-C experiment within this study and in Tüshaus et al. on differentiation day 6 relative to day 0 (n=21).

**Figure S9:**
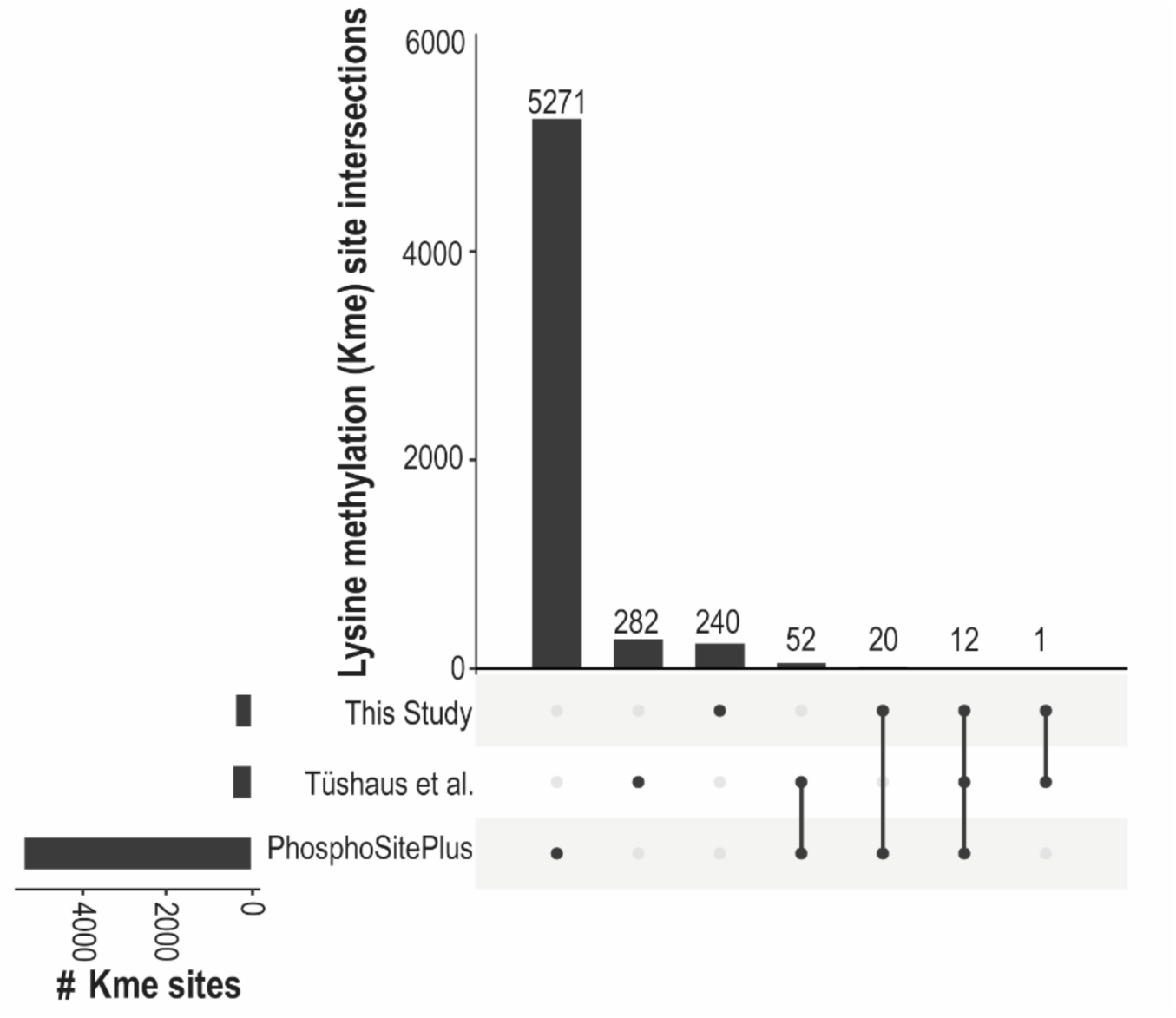
Comparison of Kme sites detected in this study and those reported in the literature. Upset plot displaying lysine methylation sites detected in this study, those reported in the PhosphoSitePlus repository, and those identified in Tüshaus et al. Horizontal bars on the left display the total number of lysine methylation sites identified. Vertical lines connecting points represent overlap of identified Kme sites.

**Figure S10:**
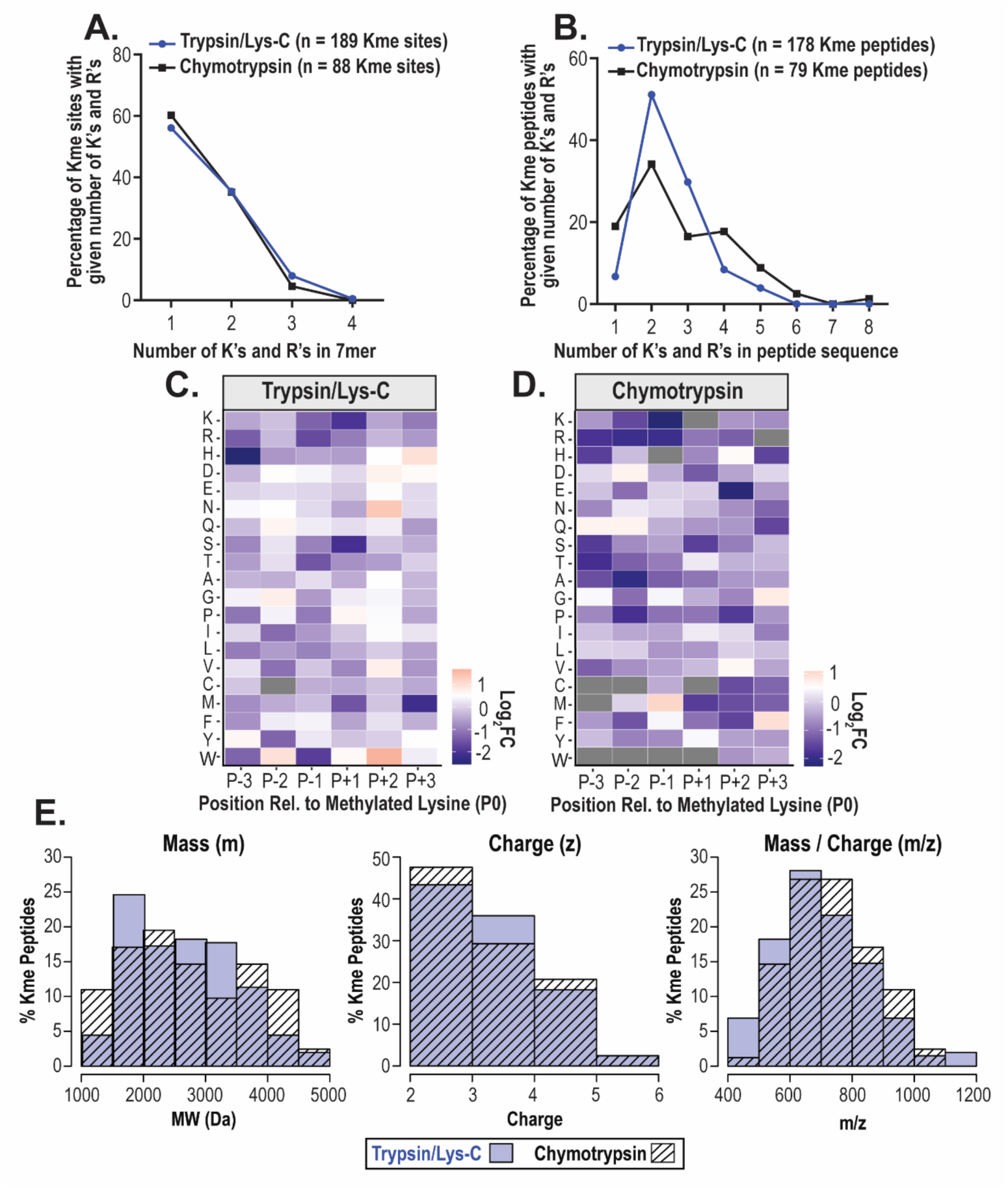
Properties of Kme sites and peptides detected in the trypsin/lys-C and chymotrypsin experiments. Lysine (K) and arginine (R) density of **(A)** 7-mer sequences surrounding methylated lysine residues and **(B)** Kme peptide sequences detected in the mass spectrometry experiments following digestion with trypsin/lys-C (blue) or chymotrypsin (black). **(C)** Trypsin/lys-C and **(D)** chymotrypsin heatmaps depicting the log_2_ ratio of the frequency of amino acids within 7-mer motifs surrounding methylated lysine residues compared to the frequency of amino acids within all lysine-centered 7-mer motifs within the human proteome. Gray squares indicate that the amino acid in that position was not present in 7-mer motifs from our study. **(E)** Histograms depicting percentages of Kme peptides detected by mass spectrometry with a given mass (m), charge (z), or mass-to-charge ratio (m/z) following digestion with trypsin/lys-C (light purple bars) or chymotrypsin (white bars with black horizontal stripes).

**Figure S11:**
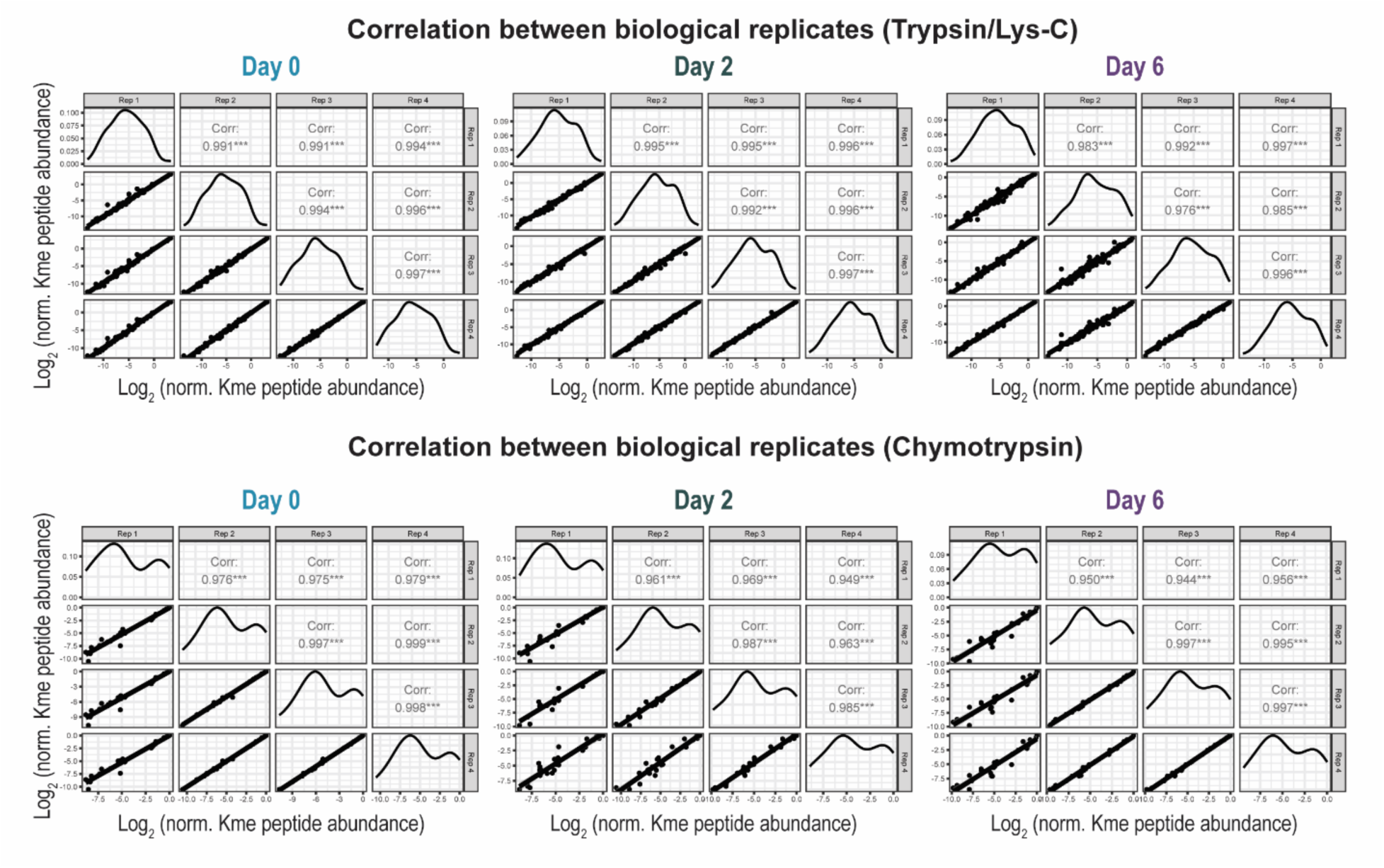
Correlation of Kme peptide abundance across biological replicates within the trypsin/lys-C and chymotrypsin experiments. Pearson correlation analysis of the average log_2_ normalized abundance of quantified Kme peptides between biological replicates within the trypsin/lys-C experiment (n=120) and within the chymotrypsin experiment (n=26).

**Figure S12:**
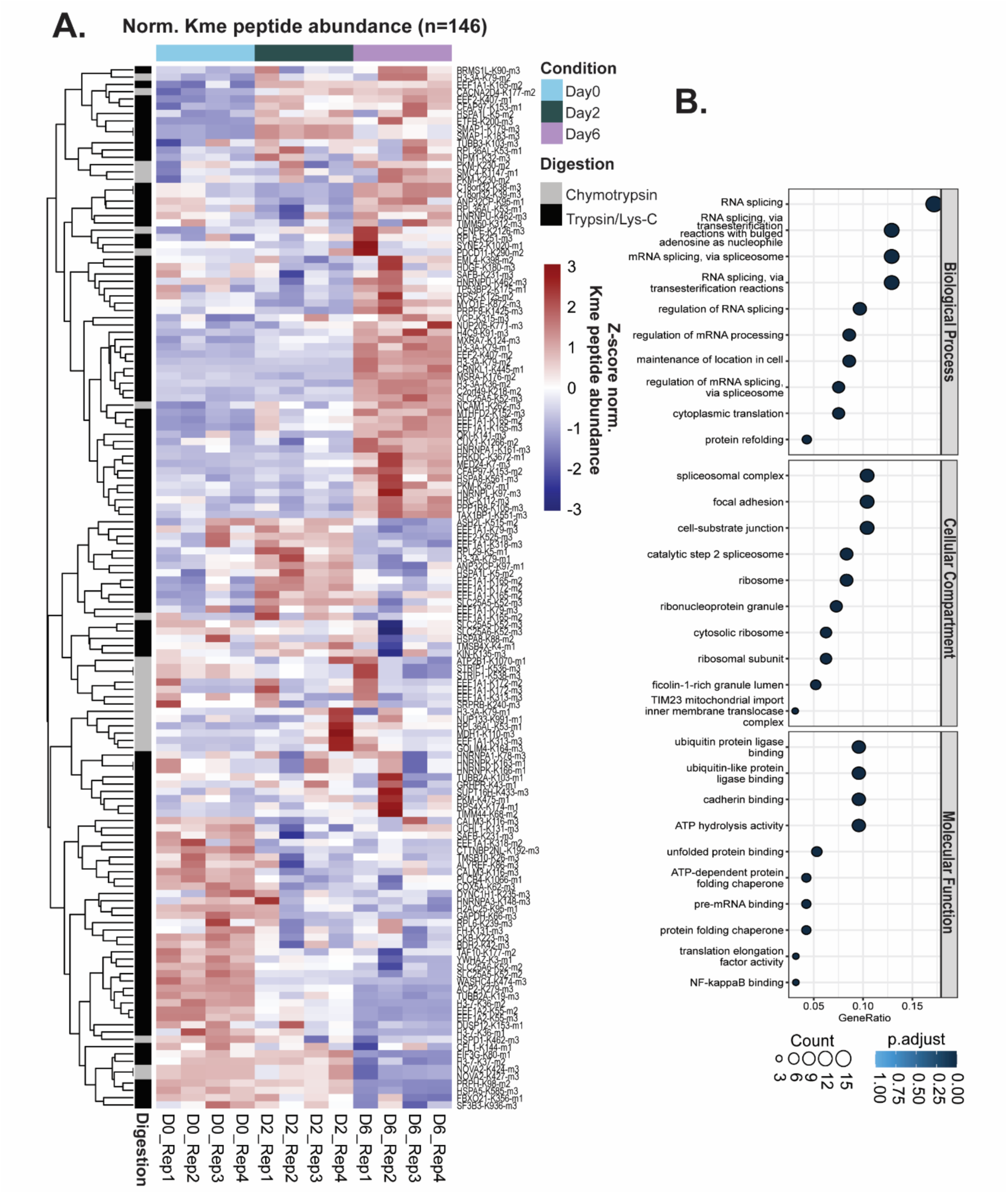
Heatmap of all Kme sites quantified in this study. **(A)** Heatmap of all Kme peptides (n=146) corresponding to 127 unique Kme sites quantified across LUHMES differentiation. Colors represent the z-score of normalized Kme peptide abundance. Rows (Kme sites) and columns (differentiation sample) are clustered by Euclidian distance. Represented to the left of the rows is the digestion strategy following which the Kme peptide was quantified. **(B)** Enriched gene ontology (GO) terms for 98 unique proteins corresponding to the Kme sites (p<0.05).

**Figure S13:**
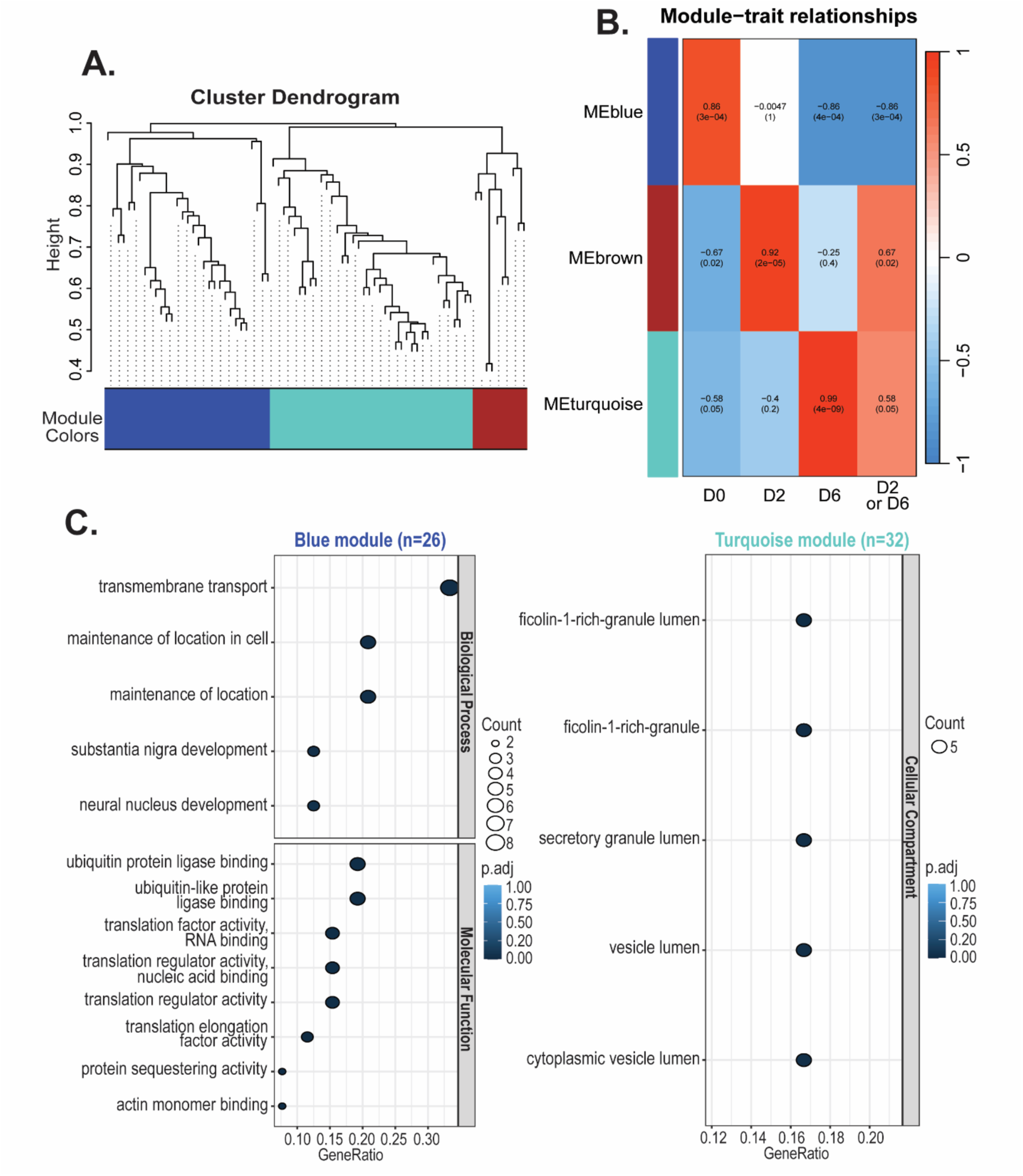
Distinct Kme peptide co-expression modules and associated functions revealed through WGCNA and GO terms analyses. **(A)** WGCNA dendrogram and **(B)** module-trait relationship correlation of Kme peptide co-expression modules identified by WGCNA analysis of differentially abundant Kme peptides. **(C)** Enriched GO terms for proteins corresponding to differentially abundant Kme peptides associated with the following modules: blue (enriched expression differentiation day 0) and turquoise (enriched expression differentiation day 6). There were no enriched biological processes, cellular compartments, or molecular functions for the brown module (enriched expression differentiation day 2).

**Table S1.**

|  | Trypsin/Lys-C |  |  | Chymotrypsin |  |  | Trypsin/Lys-C & Chymotrypsin Combined |  |  |
| --- | --- | --- | --- | --- | --- | --- | --- | --- | --- |
|  | # Kme peptides | # Kme sites | # Corr. proteins | # Kme peptides | # Kme sites | # Corr. proteins | # Kme peptides | # Kme sites | # Corr. proteins |
| Detected | 212 | 191 | 155 | 91 | 88 | 73 | 303 | 273 | 222 |
| Quantified | 120 | 108 | 84 | 26 | 24 | 18 | 146 | 127 | 98 |
| Differentially Abundant | 75 | 70 | 54 | 4 | 4 | 3 | 79 | 74 | 57 |

**Table S2.**
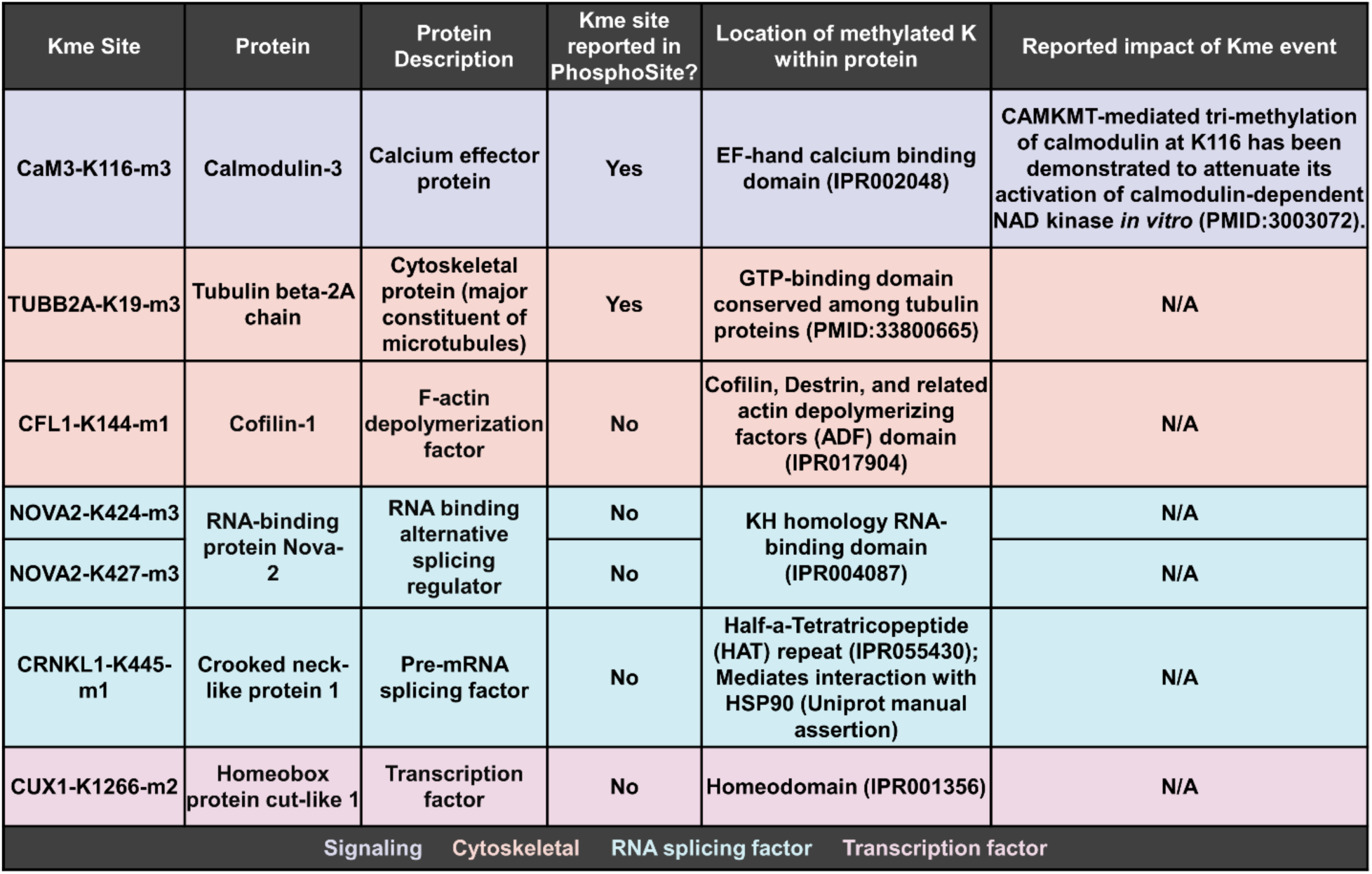

**Table S3.**
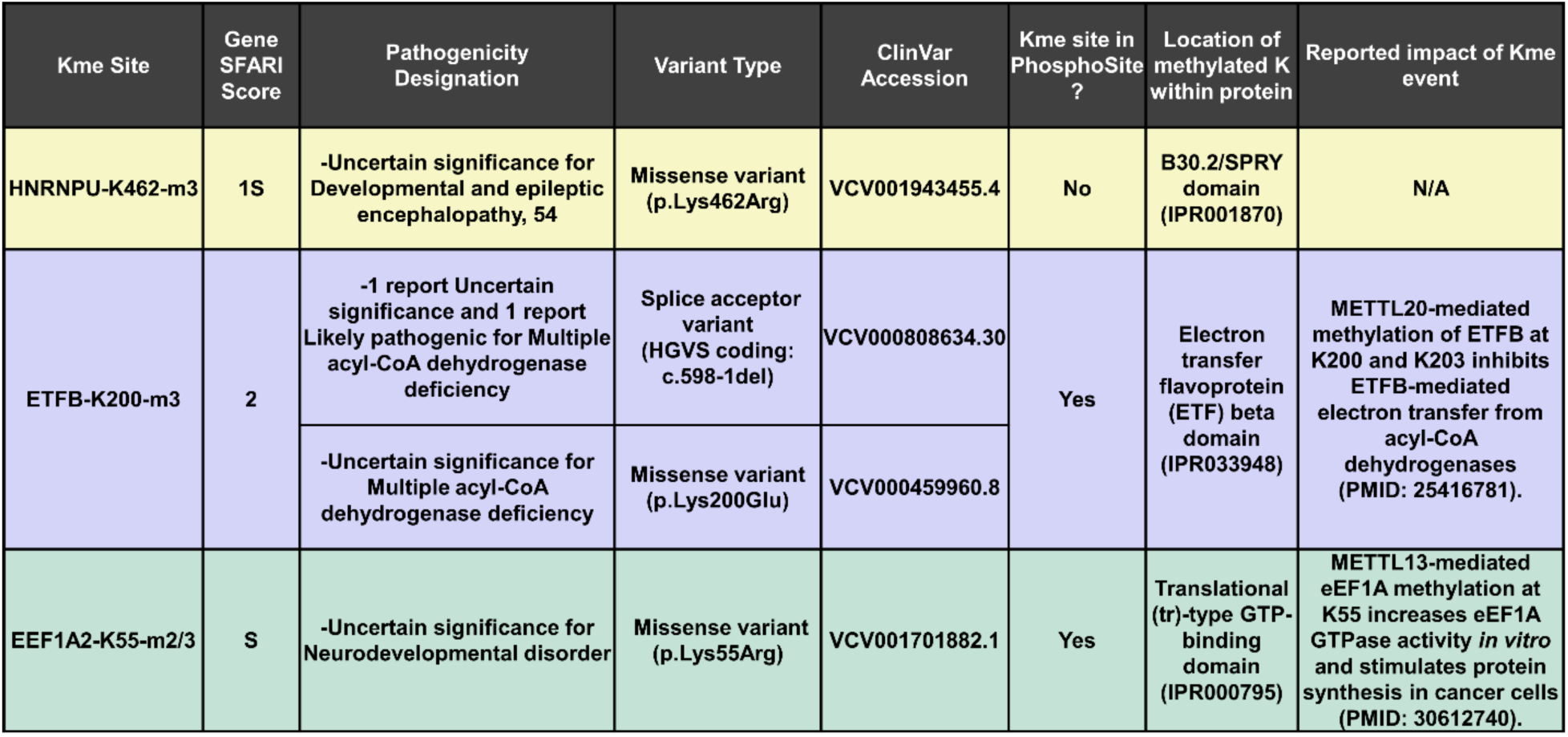

